# Oxytocin restores context-specific hyperaltruistic preference

**DOI:** 10.1101/2024.09.03.611034

**Authors:** Hong Zhang, Yinmei Ni, Jian Li

## Abstract

An intriguing advancement in recent moral decision-making research suggests that people are more willing to sacrifice monetary gains to spare others from suffering than to spare themselves, yielding the hyperaltruistic tendency. Other studies, however, indicate an opposite egoistic bias in that subjects are less willing to harm themselves for the benefits of others than for their own benefits. These results highlight the delicate inner workings of moral decision and call for a mechanistic account of hyperaltruistic preference. We investigated the boundary conditions of hyperaltruism by presenting subjects with trade-off choices combing monetary gains and painful electric shocks, or, choices combing monetary losses and shocks. We first showed in study 1 that switching the decision context from gains to losses effectively eliminated the hyperaltruistic preference and the decision context effect was associated with the altered relationship between subjects’ instrumental harm (IH) trait attitudes and their relative pain sensitivities. In the pre-registered study 2, we tested whether oxytocin, a neuropeptide linked to parochial altruism, might restore the context-dependent hyperaltruistic preference. We found that oxytocin increased subjects’ reported levels of framing the task as harming (vs. helping) others, which mediated the correlation between IH and relative pain sensitivities. Thus, the loss decision context and oxytocin diminished and restored the mediation effect of subjective harm framing, respectively. Our results help to elucidate the psychological processes underpinning the contextual specificity of hyperaltruism and carry implications in promoting prosocial interactions in our society.

## Introduction

The Chinese proverb “A virtuous man acquires wealth in an upright and just way” stresses the universal moral code of refraining from harming others for personal gain^1^. Disregard of the suffering of others is often associated with aggressive and antisocial tendencies in psychopathy^2,3^. Recent studies further developed the moral decision theory and suggested that people are willing to pay more to reduce other’s pain than their own pain, yielding a “hyperaltruistic” preference in moral dilemmas^4–7^. For example, it was shown that the hyperaltruistic preference modulated neural representations of the profit gained from harming others via the functional connectivity between the lateral prefrontal cortex, a brain area involved in moral norm violation, and profit sensitive brain regions such as the dorsal striatum^6^. However, moral norm is also context dependent: vandalism is clearly against social and moral norms yet vandalism for self-defense is more likely to be ethically and legally justified (the Doctrine of necessity). Therefore, a crucial step is to understand the boundary conditions for hyperaltruism. We set out to address this question by examining how the moral perception of decision context affects people’s hyperaltruistic prefernence^8–12^. More importantly, we also test whether the contextual effect of hyperaltruistic disposition is susceptible to oxytocin, a neuropeptide heavily implicated in social bounding and social cognition^13–15^.

Classic moral dilemmas often involve the tradeoff between personal material wellbeing and the adherence of social norm (or moral principle)^16,17^. Previous studies have shown that the moral preference can be highly context specific. For example, studies showed that people were more likely to engage in unethical behavior (lie or cheat) to avoid monetary loss than to secure monetary gain^18–20^. Other research, instead, found that it was more common for people to help others from losing money than to help them to gain money ^21,22^. However, the boundary conditions for hyperaltruism remains elusive^4–7^. Hyperaltruistic disposition was typically measured in a money-pain trade-off task where individuals’ monetary gain was pitted against the physical pain experienced by others and themselves. In turn, the hyperaltruistic preference was calculated by comparing the amounts of monetary gain subjects were willing to forgo to reduce others’ pain relative to the reduction of their own pain. Therefore, it is likely that replacing monetary gain with monetary loss in the money-pain trade-off task might bias subjects’ hyperaltruistic preference due to stronger sensitivity toward monetary loss (loss aversion) or highlighted vigilance in the face of potential loss^23–28^. Indeed, elevated attention to losses can alternate subjects’ sensitivities to the general reinforcement structure and have distinct effects on their arousal and performance^27^. Additionally, prospective losses can elicit negative emotions^29,30^, which may subsequently influence how individuals evaluate their own pain relative to others’. For example, studies have shown that negative mood can both increase pain sensitivities and decrease empathy for others, both of which yield opposing effects on hyperaltruistic preferences^31–35^. Finally, how the moral dilemma is framed can be a critical factor influencing moral decision-making^36–38^. For instance, explicitly manipulating moral frames as “harming others” (harm frame) or “not helping others” (help frame) resulted in a significant social framing effect such that subjects showed stronger prosocial preference in the harm frame compared to the help frame^25,39^. It is thus possible that people may implicitly form moral impressions (help or harm) based on the decision contexts and adjust their behavior accordingly. This way, the contextual effect on hyperaltruistic preference can be associated with individuals’ internal framing of moral contexts.

Oxytocin, a neuropeptide synthesized in the hypothalamus, has been shown to modulate a variety of emotional, cognitive and social behaviors through distributed receptors in various brain areas^40,41^. Oxytocin has been shown to play a critical role in social interactions such as maternal attachment, pair bonding, consociate attachment and aggression in a variety of animal models^42,43^. Humans are endowed with higher cognitive and affective capacities and exhibit far more complex social cognitive patterns^44^. It has been suggested that oxytocin increases prosocial behaviors such as trust and altruism by enabling individuals to overcome proximity avoidance, enhancing empathy for others’ suffering, and reducing the processing of negative stimuli (fearful or angry faces) ^45–52^. However, the evidence for whether and how oxytocin influences hyperaltruism remains scarce^53^. This may be partly because the prosocial effect of oxytocin seems to be both personality trait and decision context dependent^13,54–57^. For example, recent studies have shown that oxytocin both promotes in-group cooperation and defensive aggression toward competing out-group members^58–60^. It is therefore plausible that oxytocin might also exert a context-dependent effect on hyperaltruistic preference in a moral decision task. Specifically, oxytocin might influence the way subjects internally frame the task as benefiting from others’ pain or sacrificing self-interest to avoid others’ harm given the widely reported link between oxytocin and empathy^46,61–63^. Also, previous literature on the prosocial effect of oxytocin dovetails nicely with the utilitarian moral literature suggesting that moral behavior is closely related to people’s personality traits such as the instrumental harm attitude (IH), the degree to which people are willing to compromise their moral beliefs by inflicting harm on others to achieve better outcomes^1^.

We conducted two studies to test the above hypotheses. In study 1, we manipulated the decision context of the monetary valence (from making monetary gains to avoiding monetary losses) in a well-established money-pain trade-off task to examine the contextual specificity of the hyperaltruistic preferences (Fig. 1A). We compared subjects’ hyperaltruistic preferences in decision scenarios where options were constructed such that higher monetary gains (or smaller monetary loss in the loss context) were associated with more painful electric shocks (more pain option), and lower monetary gains (or bigger monetary loss in the loss context) were associated with less painful shocks (less pain option). We found that in line with results reported in previous studies, subjects showed clear hyperaltruistic preference in the gain context. However, shifting the decision context from gains to losses eliminated such preference. In study 2, we employed a pre-registered, placebo-controlled experimental design to examine how oxytocin might modulate the context effect of hyperaltruism (Fig. 1B). We found that oxytocin had no effect on hyperaltruistic preference in the gain context. However, oxytocin restored subjects’ hyperaltruistic preferences in the loss context.

**Fig. 1.**
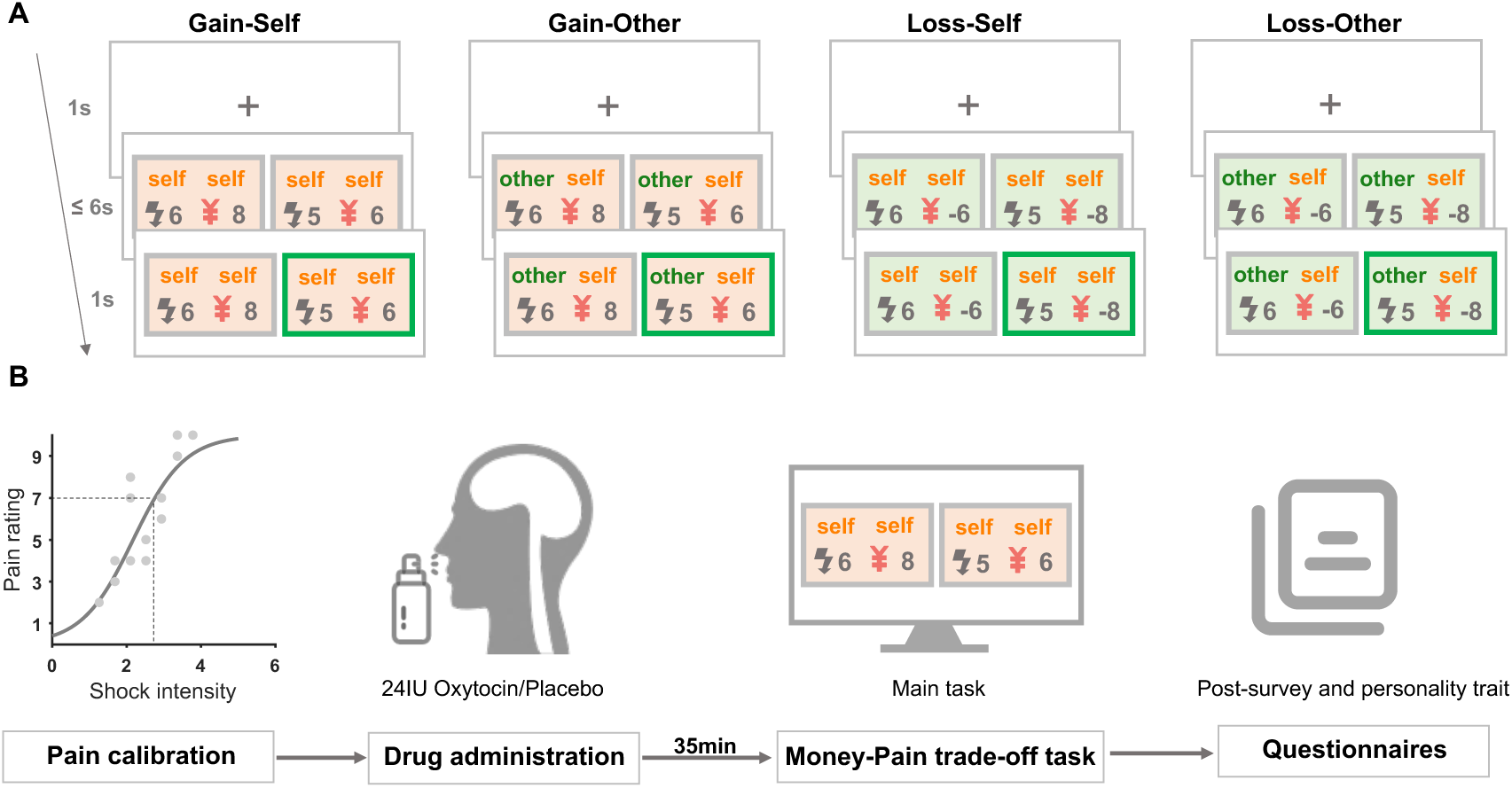
Experimental design and task. (**A**) Experimental Task. Subjects performed a money-pain trade-off task in which they were designated as deciders. Four conditions (Gain-Self, Gain-Other, Loss-Self, Loss-Other) were introduced across decision contexts (gain vs. loss) and shock-recipients (self vs. other). In each trial, subjects were asked to choose between two options with various amounts of monetary and harm consequences. The chosen option was highlighted for 1s after subjects’ decisions. (**B**) Procedures of the oxytocin study (study 2). Before the task, a pain calibration procedure was performed on each subject to determine their pain thresholds for electrical shock stimuli. Subjects were then administered with 24IU oxytocin nasal spray or placebo (saline). Thirty-five minutes later, subjects commenced the money-pain trade-off task. Finally, they filled out questionnaires including post-task surveys and assessments of personality traits.

Importantly, oxytocin administration prompted subjects to more likely perceive the task structure as harming others, which in turn mediated their hyperaltruistic preferences.

## Results

Prior to the money-pain trade-off task, we individually calibrated each subject’s pain threshold using a standard procedure^4–6^. This allowed us to tailor a moderate electric stimulus that corresponded to each subject’s subjective pain intensity. Subjects then engaged in 240 decision trials (60 trials per condition), acting as the “decider” and trading off between monetary gains or losses for themselves and the pain experienced by either themselves or an anonymous “pain receiver” (gain-self, gain-other, loss-self and loss-other, see Supplementary Fig. 9 and methods section for details).

### Loss context eliminated hyperaltruistic preference

We first tested whether the decision context (gain or loss) directly affected subjects’ hyperaltruistic tendencies by examining the proportion of trials in which subjects chose the less painful option both for themselves and for others (Fig. 1A). Since the choice sets for the self- and other-recipient are the same, subjects’ hyperaltruistic tendencies can be examined by the choice differences regarding different shock recipients (self vs. other). A larger choice difference between other- and self-recipients indicates a stronger hyperaltruistic tendencies in both the gain and loss contexts. The main effects of the shock recipient (self vs. other) and decision context (gain vs. loss) were both significant (recipient: *F*_1,79_ = 6.625, *P* = 0.012, *η*^2^ = 0.077 ; valence: *F*_1,79_ = 8.379, *P* = 0.005, *η*^2^ = 0.096), indicating that subjects were in general hyperaltruistic across decision contexts and more likely to choose the more painful option in the loss context (Fig. 2A). There was also a significant recipient × valence interaction effect (*F*_1,79_ = 33.047, *P* < 0.001, *η*^2^ = 0.295), suggesting that the loss context (vs. gain context) significantly decreased subjects’ hyperaltruistic tendencies (Fig. 2A). Further simple effect analysis confirmed that hyperaltruism only existed in the gain context (*F*_1,79_ = 16.798, *P* < 0.001, *η*^2^ = 0.175) but was eliminated in the loss context (*F*_1,79_ = 0.650, *P* = 0.423, *η*^2^ = 0.008). We also modeled subjects’ choices using an influential model where subjects’ behavior could be characterized by the harm (electric shock) aversion parameter κ, reflecting the relative weights subjects assigned to Δm and Δs, the objective difference in money and shocks between the more and less painful options, respectively (ΔV = (1- κ)Δm - κΔs Eq.1, See Methods for details)^4–6^. Higher κ indicates that higher sensitivity is assigned to Δs than Δm and vice versa. Consistent with previous literature, we found that the harm aversion parameter κ for the other- recipient was significantly greater than that of the self-recipient in the gain context^4–6^; however, such harm aversion asymmetry between self- and other-recipient disappeared in the loss context (*κ_other_* - *κ_self_* > 0 in the gain context: *t*_79_=3.869, *P* < 0.001 ; *κ_other_* and *κ_self_* showed no difference in the loss context: *t*_79_ = 0.834, *P* = 0.407; *κ_other_* - *κ_self_* showed context difference: *t*_79_ = 5.591, *P* < 0.001 Fig. 2B).

**Fig. 2.**
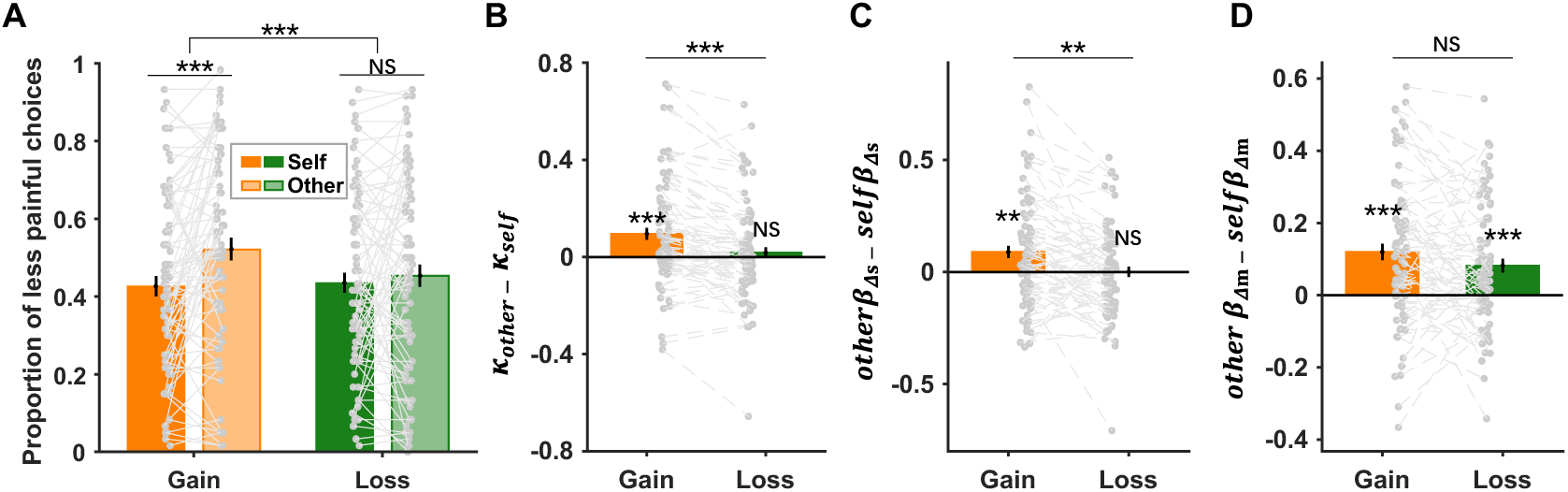
Context specific hyperaltruistic preferences. (**A**) In the gain context, subjects chose the less painful option more frequently for others than for themselves, demonstrating a hyperaltruistic preference. However, this tendency was not observed in the loss context. (**B**) The harm aversion parameter κ for others was significantly greater than that of self in the gain but not the loss context. (C) Furthermore, the relative harm sensitivity, calculated as the difference of regression coefficients of Δs in the other- and self-conditions (*otherβ_Δs_* − *selfβ_Δs_*) was significant in the gain context, but not in the loss context. (**D**) However, the relative money sensitivity, the difference of regression coefficients of Δm in the other- and self-conditions (*otherβ_Δm_* − *selfβ_Δm_*), did not show contextual specificity. Error bars represent SE across subjects. NS, not significant; ** *P* < 0.01 and *** *P* < 0.001.

To further identify the distinct effects of Δm and Δs in biasing subjects’ choice behavior, we also ran a mixed-effect logistic regression analysis to examine how the change of decision context (gain vs. loss) affected each individual’s sensitivities to money (Δm) and electric shock (Δs). Subjects’ choices were regressed against Δm and Δs and this analysis yielded significant effects on relative harm sensitivity (*other β*_Δ’_ *- self β*_Δ’_: difference of regression coefficients of Δs in the other- and self-conditions) in the gain context but not in the loss context (gain context: *t*_79_ = 3.298, *P* = 0.001; loss context: *t*_79_ = 0.015, *P* = 0.988) and significant decision context effect (*t*_79_ = 3.630, *P* = 0.001; Fig. 2C). On the contrary, decision context did not have an effect on subjects’ relative sensitivities towards monetary (Δm) difference (*t*_79_ = 1.480, *P* = 0.143), though the relative money sensitivities (*other β*_Δ+_*- self β*_Δ+_ : difference of regression coefficients of Δm in the other- and self-condition) were significant in both contexts (gain context: *t*_79_ = 5.241, *P* < 0.001; loss context: *t*_79_ = 4.355, *P* < 0.001; Fig. 2D).

These results suggest that contrary to the prediction of loss aversion, the relative money sensitivity did not change when the task structure switched from monetary gains to losses. Instead, subjects’ susceptibilities to harm might underlie the diminishment of hyperaltruism across decision contexts (also see Supplementary Fig. 1).

### Individual differences in moral preference

Recent theoretical development in moral decision-making stresses the importance of distinctive dimensions of instrumental harm and impartial beneficence in driving people’s moral preference^64,65^. The instrumental harm (IH) and impartial beneficence (IB) components of the Oxford utilitarianism scale^1^ of moral psychology were used to measure the extent to which individuals are willing to compromise their moral beliefs by breaking the rules or inflicting harm on others to achieve better outcomes (IH) and the extent of the impartial concern for the well-being of everyone (IB), respectively. We tested how hyperaltruism was related to both IH and IB across decision contexts using an exploratory multiple regression analysis. Moral preference, defined as *κ_other_* -*κ_self_*, was negatively associated with IH (*β* = -0.031 ± 0.011, *t*156 =-2.784, *P* = 0.006) but not with IB (*β* = 0.008 ± 0.016, *t*156 =0.475, *P* = 0.636) across gain and loss contexts, reflecting a general connection between moral preference and IH (Fig. 3A & B). Note that the interaction effects of valence × IH and valence × IB were not significant (valence × IH effect: *β* = 0.016 ± 0.022, *t*152 = 0.726, *P* = 0.469, Fig. 3A; valence × IB effect: *β* = -0.004 ± 0.031, *t*152 = -0.115, *P* = 0.908, Fig. 3B).

**Fig. 3.**
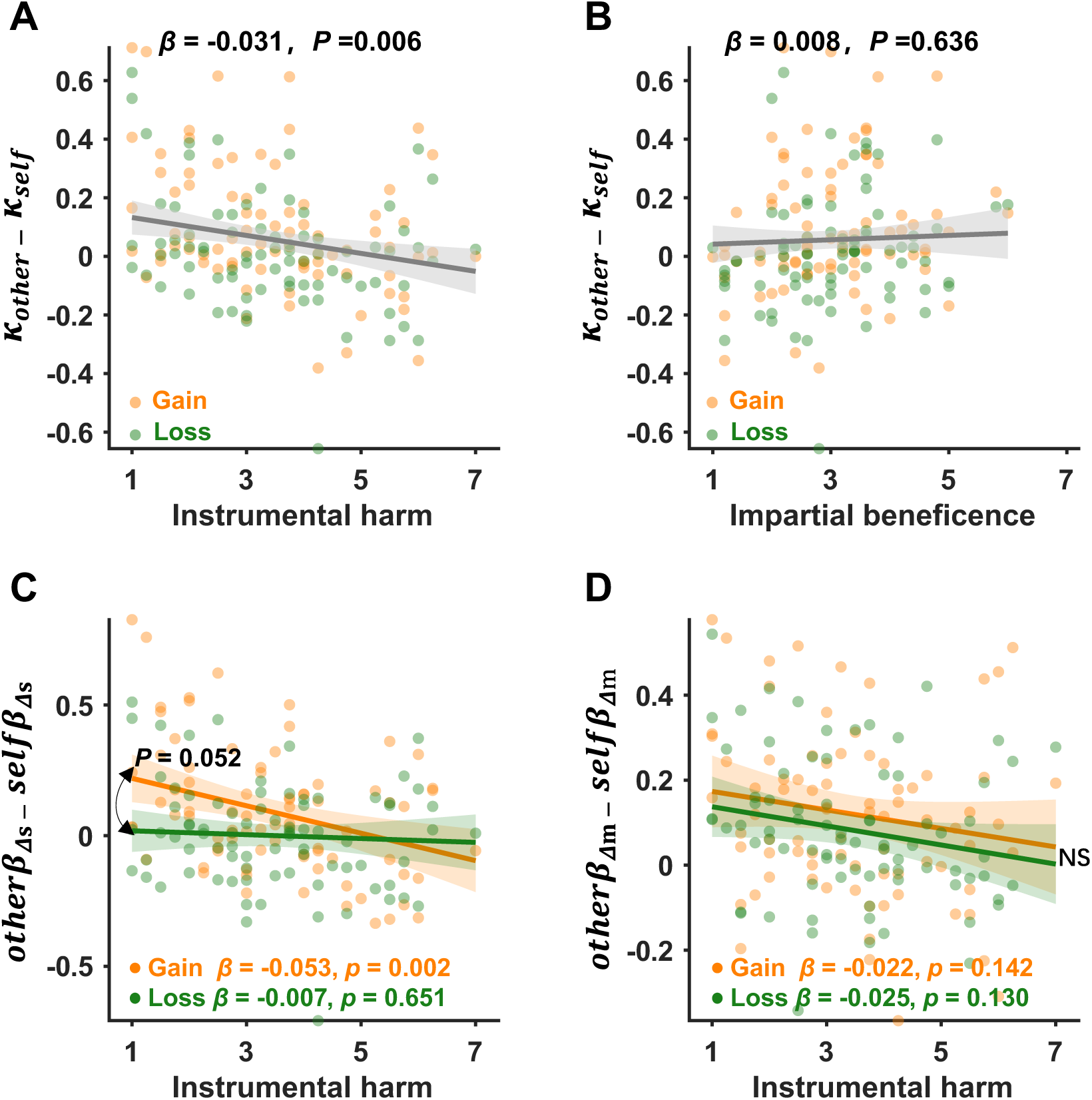
Individual difference in moral preferences. (**A-B**) Hyperaltruism (*κ*_$%&’(_ − *κ*_)’*+_) was negatively associated with instrumental harm (IH) attitudes but showed no significant relationship with impartial beneficence (IB). (**C**) The correlation between IH and subjects’ relative harm sensitivities (*otherβ_Δs_* − *selfβ_Δs_*) was marginally different between the gain and loss contexts. (D) However, the correlation between IH and subjects’ monetary sensitivities (*otherβ_Δm_* − *selfβ_Δm_*) showed no context difference. The regression coefficients in Figure A, C & D showed the relationship between IH and moral behavior, controlling for empathic concern (EC) and impartial beneficence (IB), while the coefficient in Figure B showed the relationship between IB and moral preference, controlling for EC and IH. NS, not significant.

Interestingly, by examining the separate contributions of Δm and Δs to moral preferences, we found that the decision context modulated the relationship between IH and subjects’ relative harm sensitivities (gain context: *β* = -0.053 ± 0.016, *t*152 =-3.224, *P* = 0.002; loss context: *β* = -0.007 ± 0.016, *t*152 =-0.453, *P* = 0.651; valence × IH: *β* = 0.045 ±0.023, *t*152= 1.959, *P* = 0.052 ; Fig. 3C), yet it did not modulate the relationship between IB and subjects’ relative harm sensitivities (gain context: *β* = 0.019 ± 0.023, *t*152 =0.804, *P* = 0.423; loss context: *β* = 0.010 ± 0.023, *t*152 = 0.425, *P* = 0.672; valence × IB: *β*=-0.009 ± 0.033, *t*152 = -0.268, *P* = 0.789 ; also see Supplementary Fig. 3A). However, subjects’ monetary sensitivities were not related with IH (gain context: *β* = -0.022 ±0.015, *t*152 =-1.476, *P* = 0.142; loss context: *β* = -0.023 ±0.015, *t*152= -1.523, *P* = 0.130; valence × IH: *β* = -0.001 ± 0.021, *t*152 = -0.034, *P* = 0.973 ; Fig. 3D), nor with IB (gain context: *β* = -0.008 ± 0.021, *t*152 = -0.369, *P* = 0.713 ; loss context: *β* = -0.009 ± 0.021, *t*152 = -0.414, *P* = 0.679 ; valence × IB: *β* = -0.001 ± 0.030, *t*152 = -0.032, *P* = 0.974 ; Supplementary Fig. 3B; also see Supplementary Fig. 2 and Table S1). These results suggest that the context dependent moral preference may be specifically due to the context modulation of subjects’ relative harm sensitivities, accompanied by the altered correlation between relative harm sensitivities and IH across decision contexts. Furthermore, our results are consistent with the claim that profiting from inflicting pains on another person (IH) is inherently deemed immoral^1^. Hyperaltruistic preference, therefore, is likely to be associated with subjects’ IH dispositions.

### Oxytocin restored hyperaltruistic preferences in the loss context

To probe how oxytocin might alter subjects’ moral preference, we conducted the preregistered study 2 where a separate cohort of subjects performed the same task while undergoing the placebo/oxytocin administration in a with-subject design (N = 46, see Methods for experimental details). First, in the placebo session of study 2, we replicated the major findings of study 1 (Fig. 2A), demonstrating that hyperaltruistic preference was not significant in the loss context compared to the gain context (Placebo: valence × recipient interaction: *F*_1,45_ = 9.103, *P* = 0.004, *η*^2^ = 0.168; simple effect: gain context *F*_1,45_ = 9.356, *P* = 0.004, *η*^2^ = 0.172; loss context *F*_1,45_ = 0.005, *P* = 0.946, *η*^2^ < 0.001; Fig. 4A). The main effect of treatment (placebo vs. oxytocin) was significant (*F*_1,45_ = 41.961, *P* < 0.001, *η*^2^ = 0.483), indicating that the oxytocin treatment prompted subjects to select the more painful option more often. Interestingly, the treatment × recipient × valence interaction effect was also significant (*F*_1,45_ = 6.309, *P* = 0.016, *η*^2^ = 0.123), suggesting that the recipient × valence interaction might be different due to the placebo and oxytocin treatments. Indeed, with the administration of oxytocin, the hyperaltruistic preference was restored in the loss context, without affecting the hyperaltruistic pattern in the gain context (oxytocin: valence × recipient: *F*_1,45_ = 0.480, *P* = 0.492, *η*^2^ = 0.011; simple effect: gain context: *F*_1,45_ = 25.364, *P* < 0.001, *η*^2^ = 0.360; loss context: *F*_1,45_ = 24.408, *P* < 0.001, *η*^2^ = 0.352; Fig. 4A). These results together highlight that oxytocin might play an important valence-dependent modulatory role in restoring hyperaltruistic moral preference. Our modeling analysis revealed similar results: there was a significant treatment × valence interaction effect (*F*_1,45_ = 6.349, *P* = 0.015, *η*^2^ = 0.124) on subjects’ moral preference (defined as *κ_other_* - *κ_self_*, Fig. 4B). With the administration of oxytocin, however, the diminished hyperaltruistic preference in the loss context (placebo: *F*_1,45_ = 1.295, *P* = 0.261, *η*^2^ =0.028) was successfully restored (oxytocin: *F*_1,45_=17.990, *P* < 0.001, *η*^2^ = 0.286, simple effect analysis, Fig. 4B). Additionally, the model-based moral preference was correlated with subjects’ IH reports (but not IB) across subjects in both the placebo and oxytocin treatments (Supplementary Fig. 4 & 5, also see Table S2), replicating the results we obtained in study 1(Fig. 3A & B).

**Fig. 4.**
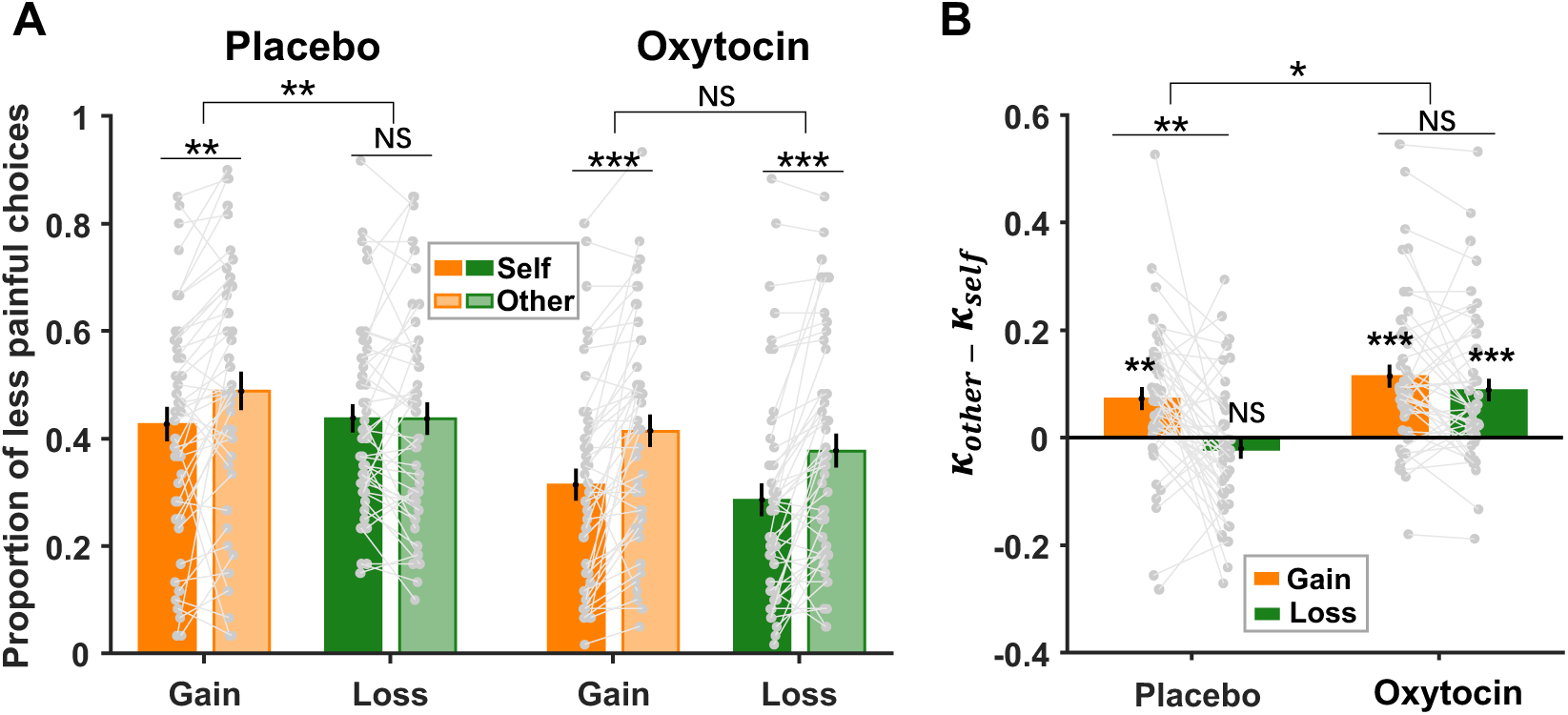
Oxytocin significantly promoted hyperaltruistic preference in the loss context. (**A**) In contrast to the placebo condition, oxytocin administration elicited hyperaltruistic behavior (a greater tendency of choosing less painful options in other condition than self-condition) in the loss context compared to the gain context. (**B**) Model-based hyperaltruistic parameter (*κ*_$%&’(_ − κ_)’*+_) showed similar patterns: Hyperaltruistic tendency was reduced by the loss context in the placebo session but restored in the oxytocin session. Error bars represent SE across subjects. NS, not significant; * *P* < 0.05, ** *P* < 0.01 and *** *P* < 0.001.

### Hyperaltruistic preference and increased harm sensitivities with Oxytocin administration

In study 1, we showed that the lack of hyperaltruism in the loss context was specifically related to the diminished relative harm sensitivities. In study 2, we also ran a mixed-effect logistic regression of subjects’ choices against continuous independent variables including Δm, Δs and categorical variables including treatment (placebo vs. oxytocin), harm/shock recipient (self vs. other) and decision context valence (gain vs. loss; see methods for details). This regression analysis yielded a significant Δs × treatment × recipient × valence interaction effect (*β* = 0.125 ± 0.053, *P* = 0.018, 95% CI = [0.022, 0.228]; Supplementary Fig.6 & 8), indicating the relative harm sensitivities (*other β*_Δ’_ − *self β*_Δ’_) was modulated by the specific treatment and decision combinations. Indeed, repeated ANONA analysis confirmed that in the placebo session, as in study 1, the relative harm sensitivities decreased in the loss context (relative to the gain context, simple effect: *F*_1,45_ =11.521, *P* = 0.001, *η*^2^ = 0.204; Fig. 5A); however, with the administration of oxytocin, there was no significant difference of the relative harm sensitivities between the gain and loss contexts (simple effect: *F*_1,45_ = 0.131, *P* = 0.719, *η*^2^ = 0.001; Fig. 5A). It is worth noting that oxytocin did not alter the gain/loss relative money sensitivity difference (valence × treatment interaction: *F*_1,45_ = 1.933, *P* = 0.171, *η*^2^ = 0.041; Fig. 5B), which corresponded to a non-significant Δm × treatment × recipient × valence interaction effect (*β* = -0.052 ± 0.100, *P* = 0.601, 95% CI = [-0.249, 0.144]; Supplementary Fig. 6). Furthermore, in the placebo session, the decision context significantly modulated the relationship between the relative harm sensitivity and IH (gain context: *β* = -0.071 ± 0.024, *t*_84_ = -3.005, *P* = 0.004; loss context: *β* = 0.006 ± 0.024, *t*_84_ = -0.243, *P* = 0.809; valence × IH: *β* = 0.077 ± 0.033, *t*_84_ =2.297, *P* = 0.024; Fig. 5C). However, under the treatment of oxytocin, the valence × IH interaction became non-significant (*β* = -0.006 ± 0.021, *t*_84_ =-0.272, *P* = 0.786; Fig. 5D), despite the significant negative relationship between the relative harm sensitivities and IH in both decision contexts (gain context: *β* = -0.037 ± 0.015, *t*_84_ = -2.458, *P* = 0.016; loss context: *β* = -0.042 ± 0.015, *t*_84_ = -2.843, *P* = 0.006; Fig. 5D) (also see Supplementary Fig. 7 and Table S2). The effect size (Cohen’s *f* ²) for this exploratory analysis was calculated to be 0.491 and 0.379 for the placebo and oxytocin conditions, respectively. The post hoc power analysis with a significance level of α = 0.05, 7 regressors (IH, IB, EC, decision context, IH×context, IB×context, and EC×context), and sample size of N = 46 yielded achieved power of 0.910 (placebo treatment) and 0.808 (oxytocin treatment). These findings further highlighted the influence of decision context on hyperaltruistic moral preference and stressed the importance of oxytocin in restoring the correlation between the relative harm sensitivity and the dispositional personality traits such as IH.

**Fig. 5.**
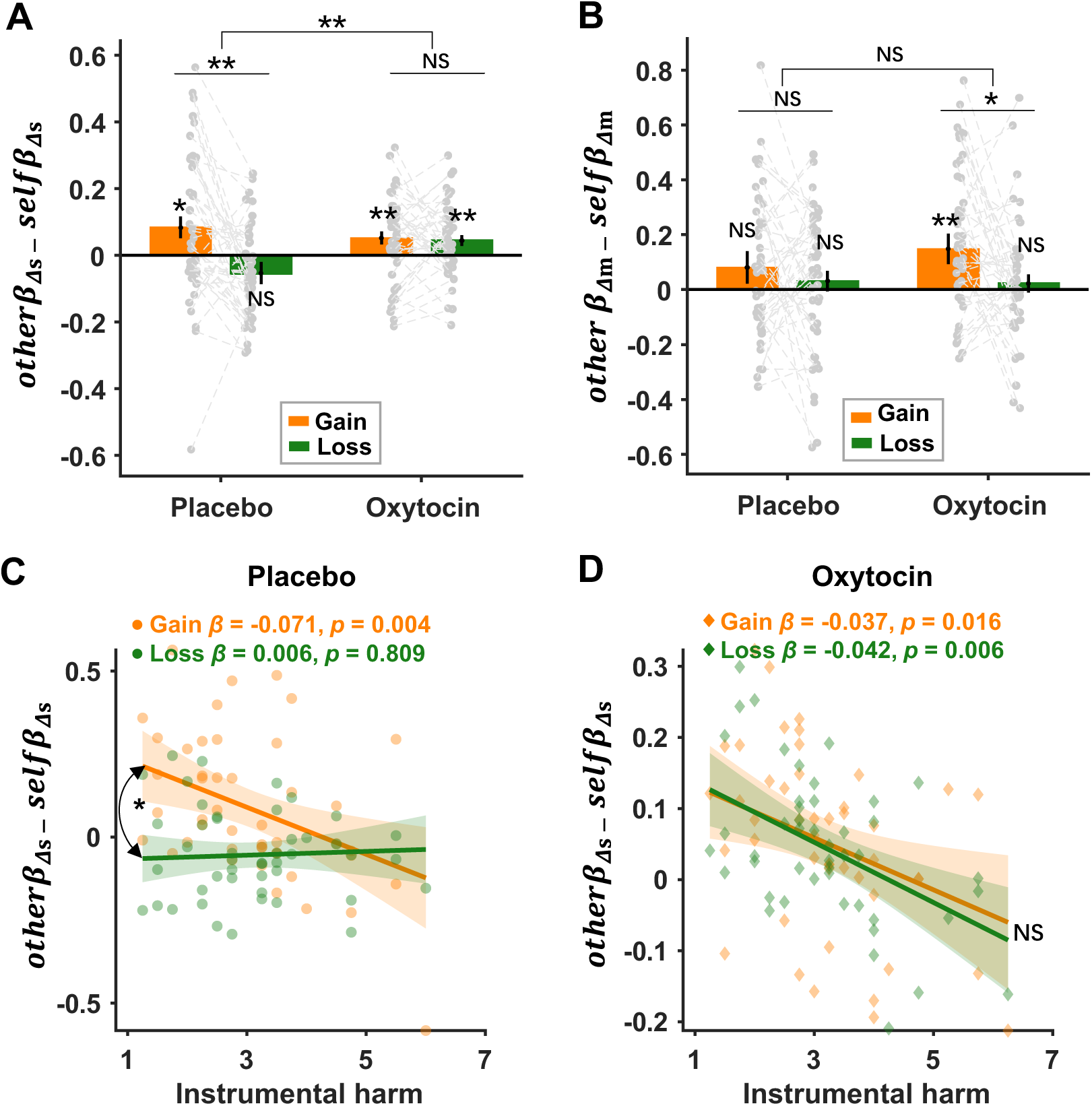
Oxytocin effect on relative harm sensitivities. (**A**) Oxytocin significantly modulated the context-specificity of relative harm sensitivity (*otherβ_Δs_* − *selfβ_Δs_*). (**B**) Oxytocin did not influence the contextual differences of the relative monetary sensitivities (*otherβ_Δm_* − *selfβ_Δm_*). (**C-D**) The decision context modulated the correlation between instrumental harm (IH) and subjects’ harm sensitivities in the placebo session (**C**), whereas oxytocin eliminated the contextual modulation (**D**). Error bars represent SE across subjects. The regression coefficients in Figure C & D showed the association between IH and relative harm sensitivities, controlling for empathic concern and impartial beneficence. NS, not significant; * *P* < 0.05, ** *P* < 0.01.

### Oxytocin eliminated the modulation effect of decision context on hyperaltruism

We reasoned that the decision context effect of hyperaltruistic preferences might be rooted in how our subjects internally framed the task as “harming” (inflicting pain on the receiver to increase monetary gain/avoid monetary loss) or “helping” others (sacrificing greater monetary gain/accepting greater monetary loss to alleviate the receiver’s pain). To test this hypothesis, in study 2 we asked subjects to subjectively report whether they perceived the task as “harmful” or “helpful” (see Methods section for more details) in both placebo and oxytocin sessions (gain and loss contexts for each session). The non-parametric Friedman tests on subjects’ harm framing report showed a significant difference in gain and loss contexts under placebo condition (*χ*^2^ = 2.827, *P* = 0.028, Bonferroni correction; Fig. 6A), suggesting that subjects were more likely to perceive the task as harming others in the gain context. In addition, the administration of oxytocin increased subjects’ perception of causing harm to others in the loss context (*χ*^2^ = 2.665, *P* = 0.046, Bonferroni correction) and removed contextual disparities in harm perception (valence × treatment interaction: *χ*^2^ = 7.410, *P* = 0.006) (Fig. 6A), suggesting that the oxytocin administration successfully erased the framing difference between the gain and loss decision contexts.

**Fig. 6.**
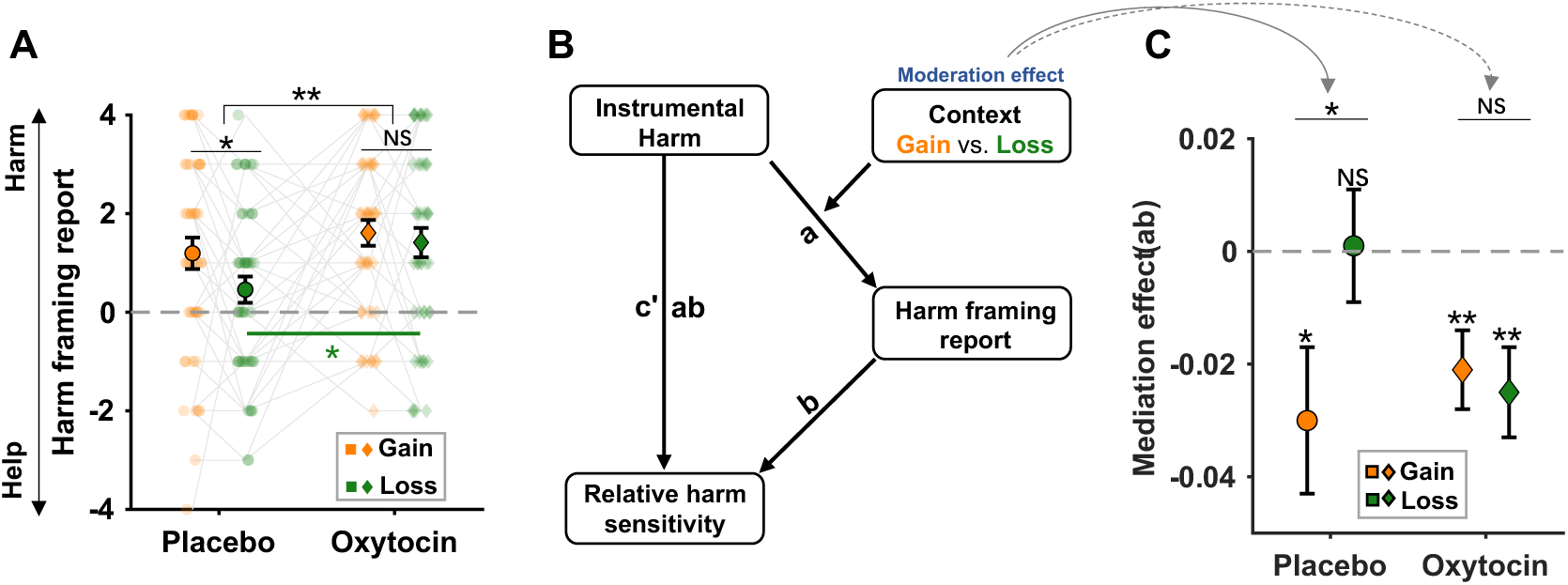
Oxytocin modulated the contextual influence on hyperaltruistic behaviors. (**A**) Monetary loss (relative to gain) significantly reduced subjects’ perception of harm framing in the task. Oxytocin augmented harm framing perception, particularly in the loss context, effectively removing the contextual specificity of harm framing perception. (**B**) The conceptual diagram of the moderated mediation model. We assume that the perceived harm framing mediates the relationship between instrumental harm and relative harm sensitivity, with decision context moderating the mediation effect. (**C**) The moderating effect of decision context was significant under placebo condition. However, oxytocin obliterated the contextual moderation effect by reinstating the mediating role of harm perception in the loss context. Error bars represent SE across subjects. NS, not significant; * *P* < 0.05, ** *P* < 0.01.

We further hypothesized that the correlations between instrumental harm (IH) personality traits and subjects’ differential harm sensitivities (*otherβ*_,’_ − *selfβ*_,’_, Fig. 5C & D) might be mediated by how strongly subjects perceived the decision task as harming or helping others. A moderated mediation analysis was thus conducted to directly test this hypothesis. In the moderated mediation model, the effects of decision context can be expressed as the moderation effects on mediation (Fig. 6B). As we expected, in the placebo session, the moderating effect of decision context was significant (placebo: the differential mediation effect in gain and loss contexts: Δab = -0.036, *P* = 0.036, 95%CI = [-0.073, -0.003] ; Fig. 6C). Specifically, the mediation effect of perceived harm was significant in the gain context (ab = -0.030, *P* = 0.021, 95%CI = [-0.056, -0.006]) but not in the loss context (ab = 0.001, *P* = 0.915, 95%CI = [-0.019,0.019]). However, oxytocin eliminated the contextual moderation effect (oxytocin: the differential mediation effect in gain and loss contexts: Δab = -0.006, *P* = 0.731, 95%CI = [-0.044, 0.033]; Fig. 6C) by reinstating the mediation role of harm perception in the loss context (ab = -0.025, *P* = 0.002, 95%CI = [-0.042, -0.011]) (also see Table S3). These results indicated that subjects’ perception of the task structure as helping or harming others mediated the correlation between personality trait IH and subjects’ hyperaltruistic preference. Switching the decision context from gains to losses directly modulated subjects’ moral perception of the task and nullified hyperaltruism. Finally, oxytocin administration restored participants’ hyperaltruistic preference by reinstating the mediation role of perceived harm report on the correlation between IH and hyperaltruism across gain and loss decision contexts.

## Discussion

Utilitarian moral decision-making studies often involve moral dilemmas where utilitarian gains are traded against moral norms such as no deception or no infliction of suffering on others^1,16,66^. Recent studies extended this line of research by comparing subjects’ sensitivities toward others’ suffering relative to their own suffering and revealed an intriguing hyperaltruistic preference: people were willing to forgo more monetary gain to reduce others’ pain than their own pain^4–7^. In two experiments, we replicated findings in the previous literature (Figs.1, 4). More importantly, we further showed that hyperaltruistic preference was susceptible to the corresponding decision context. Replacing the trade-off between monetary gain and electric shocks (gain context) with arbitration between monetary loss and electric shocks (loss context) eliminated subjects’ hyperaltruistic preference that would otherwise have been observed. Importantly, oxytocin restored the hyperaltruistic preference in the loss context by biasing subjects to more likely perceive the decision context as harming others for self-interest. Finally, moral framing, or how subjects perceive the decision context as helping or harming others, mediated the association between the instrumental harm (IH) personality trait and subjects’ sensitivities to others’ suffering relative to their own suffering. Therefore, we demonstrated that both the hyperaltruistic preference and the effects of oxytocin were context-dependent and provided a mechanistic account and the boundary condition for hyperaltruistic disposition.

Recent development in utilitarian moral psychology highlights two independent dimensions that collectively shape people’s moral preference: attitudes toward instrumental harm (IH), the suffering of other individuals to achieve greater good, and toward impartial beneficence (IB), an impartial concern for the well-being of everyone^1^. In our experiments, we found that subjects’ IH attitude was negatively associated with their hyperaltruistic preference (Fig. 3A). However, there was no significant relationship between IB attitudes and hyperaltruism (Fig. 3B), suggesting the money- pain trade-off task might be better suited to study the specific relationship between IH attitude and moral preference^16^. Further analyses decomposing subjects’ hyperaltruistic preference into their relative sensitivities towards shock difference (Δs) and money difference (Δm) suggested that the decision context (loss vs. gain) altered the relationship between IH and Δs relative sensitivity (Figs. 3C, 5C) but not the association between IH and Δm sensitivity (Fig. 3D). These results confirm that how subjects evaluate others’ suffering relative to their own suffering underlies the relationship between IH and hyperaltruistic preferences.

It might be argued that the decision context effect on hyperaltruistic preference was due to loss aversion, a phenomenon that people weigh prospective loss more prominently than monetary gain^23^. Our results, however, showed that the relative evaluation of Δm did not differ significantly across decision contexts (gain vs. loss, Figs. 2D & 5B). Furthermore, there was no significant interaction effect of IH and decision contexts on the relative evaluation of Δm (Fig. 3D, supplementary Fig. 7A), excluding the role of loss aversion in mediating the context effect on hyperaltruistic preference. From a theoretical point of view, loss aversion only proportionally shifts harm aversion parameter κ (representing the relative importance of Δm and Δs) in the self- and other- conditions, respectively, and will not change the direction of hyperaltruistic preference (*κ_other_* - *κ_self_*).

Instead, we propose that the moral framing of the decision context as helping or harming others may drive subjects’ hyperaltruistic preference. Indeed, we showed that subjects were less likely to perceive the task as harming others for monetary benefit when the decision context switched from gain to loss condition (Fig. 6A). In addition, subjects’ harm framing reports fully mediated the correlation between IH and the relative harm sensitivity in the gain conditions (Figs. 5C & 6C). Decision context exerted its effect by modulating the mediation effect of harm framing report such that the loss context eliminated the correlation between the relative harm sensitivity and IH (Fig. 5C). Our results are consistent with a recent study where the exogenous moral framing of the task (harming or helping others) significantly biased subjects’ prosocial behavior^25,39^. Subjects behaved more prosocially under the harming frame compared to the helping frame. In our experiments, the decision contexts prompted different endogenous moral framing of the task such that more harm framing led to higher hyperaltruism levels. These results taken together suggest that moral framing may be a critical factor influencing subjects’ prosocial preference. Another recent study reported that human subjects can be both egoistic and hyperaltruistic in moral decisions^7^. While the hyperaltruistic preference was elicited by comparing how subjects evaluated others’ suffering to their own suffering in the standard monetary gain-pain trade-off task, egoistic tendencies emerged when subjects had to decide whether to harm themselves for others’ benefit (compared to harming themselves for their own benefit). These ostensibly puzzling results can be reconciled under the framework of internal moral framing of the task. In order for participants to express moral or prosocial choices, they have to identify how salient a moral code is in place. Internally framing the task as benefiting from others’ suffering would be more likely to discourage people from actions they would have otherwise taken if the task is perceived as sacrificing self interest in order to help others. The personality trait instrument harm (IH) attitudes tap into subjects’ relative harm sensitivities and the correlation between IH and relative harm sensitivity is modulated by the decision context.

In our pre-registered study 2, we directly tested the potential mediating role of moral framing and found that the administration of oxytocin significantly increased subjects’ moral framing of the decision context as harming others in both the gain and loss contexts (Fig. 6A). It is worth noting that oxytocin administration did not affect subjects’ IH disposition (Supplementary Fig. 4B). Therefore, oxytocin effectively nullify the modulation effects of decision context and preserved the correlation between the relative harm sensitivity and IH in both decision contexts (Figs. 5D, 6C). Oxytocin has long been featured in the general approach-avoidance hypothesis^67^, which proposes that oxytocin attenuates people’s avoidance of negative social or nonsocial stimuli^49,67–69^. Other studies conducted on both animals and humans suggest that oxytocin increases pain tolerance and attenuates acute pain experience ^70–72^. However, hyperaltruistic preference is tied to the differential evaluation of other’s suffering relative to subjects’ own suffering and thus a general approach-avoidance theory of oxytocin might not be capable to account for the decision context specific effect. Recently, oxytocin has been shown to play a significant role in regulating parochial altruism by promoting both in- group cooperation and out-group aggression ^58–60^. Our results corroborated with this line of research and highlighted the role of oxytocin in modulating the moral perception of the decision environment. Importantly, we demonstrated that moral perception of the task structure mediated the correlation between subjects’ personality attitudes towards harming others (IH) and their relative harm sensitivities which directly contributed to the emergence of hyperaltruistic preference. Through this lens, decision-context can be viewed as the implicit proxy of the mediator (task moral perception). As oxytocin increases the task perception as more harming others for self-interest (Fig. 6A), the correlation between IH and the relative harm sensitivities is restored.

In summary, in two studies, we demonstrate subjects’ hyperaltruistic preference is closely associated with the decision context. Subjects’ internal moral framing of the decision context mediates the correlation between personality traits such as instrumental harm (IH) attitude and the relative harm sensitivity and the decision context exerts a modulation effect on the mediation effect. Intranasal oxytocin administration drives subjects to be more harm frame oriented and abolishes the modulation effects of the decision contexts. Therefore, our study provides valuable insights into the psychological and cognitive mechanisms underlying hyperaltruistic behavior and highlights the crucial role of oxytocin in shaping moral decision-making. Finally, our findings carry significant implications for future research on moral behavior and its impairment in specific crisis contexts.

## Materials and Methods

### Participants

All subjects were students recruited through online university platform with written informed consent. Subjects were free of historic or current neurological or psychological disorders and with corrected normal vision. This study was approved by the institutional review board of the school of psychological and cognitive sciences at Peking University.

We conducted a power analysis (G*Power 3.1) to determine the number of subjects sufficient to detect a reliable hyperaltruistic effect reported in the previous literatures^4–7,73^. Based on the small to medium effect size (Cohen’s d = 0.2), 75 subjects were needed to detect a significant effect (α = 0.05, β = 0.9, two-by-two within ANOVA) for study 1. We ended up recruiting 83 right-handed subjects for study 1. Two subjects were excluded from data analysis due to their exclusive selection of the same option (more or less painful) across all trials, and 1 subject did not complete the experiment, leaving a total of 80 subjects (40 males, mean age = 21.38 ± 2.67 years). Study 2 (the oxytocin study) was specifically designed to test the oxytocin effect on hyperaltruism and was preregistered (https://osf.io/fhwa9) via the Open Science Framework. Due to the potential confounds of the oxytocin effects, we only recruited male subjects for Study 2^74,75^. It should be noted that in preregistration we originally planned to recruit 60 male subjects for Study 2 but ended up recruiting 46 male subjects (mean age = 21.74 ± 2.33 years) based on the sample size reported in previous oxytocin studies^57,69^. Additionally, a power analysis suggested that the sample size > 44 should be enough to detect a small to median effect size of oxytocin (Cohen’s d = 0.21, α = 0.05, β = 0.8) using a 2 × 2 × 2 within-subject design^76^. All the subjects were instructed to avoid taking caffeine, cigarettes, and alcohol 24 hours before the experiment and to refrain from eating or drinking two hours before the experiment.

### Experimental Procedures

Each time, two subjects (a pair) arrived with a 5-minute interval and entered into separate testing rooms without seeing each other to ensure complete anonymity. After signing the informed consent, subjects completed the pain calibration procedure (described below) in separate rooms. We then followed the protocol developed in the previous literature to assign experimental roles to both subjects before the main task^4^. Briefly, each subject was told that there was a second subject in the next testing room (whom they would not meet in person). Subjects were instructed that a random coin toss would decide their roles as the decider or receiver in the upcoming money-pain tradeoff task. Unbeknownst to the subjects, both of them were assigned to the role of deciders.

For study 2, the experimental procedures were similar to those in study 1. In addition, both subjects were administrated with either oxytocin or placebo in two sessions (session 1 & 2) of 5∼7 days apart in a randomized order, yielding 11 pairs (22 subjects) having the oxytocin session first and the other 12 pairs (24 subjects) taking the placebo session first. Also, once the subjects’ task roles (deciders) were announced in session 1, their roles remained in session 2.

### Pain calibration

We adopted a standard pain titration paradigm that has been widely used in previous literature ^4,77^. Electric shocks were generated with a Digitimer DS5 electric stimulator (Digitimer, UK) and applied to the inner side of the subject’s left wrist via two electrodes. By slowly increasing or decreasing the electric shock intensities, we asked subjects to rate their pain experience on a 11-point scale ranging from 0 (no pain at all) to 10 (intolerable) until a rating of 10 was reached (subject’s maximum tolerance threshold). Next, we generated shocks ranging from 30% to 90% of the subjects’ maximum threshold in 10% increments. Each shock intensity was delivered three times. Subjects in total received 21 shocks in randomized order and gave pain ratings on the 11-point scale. For each subject, we fitted a sigmoid function to subjects’ subjective pain ratings and chose the current intensity that corresponded to each subject’s subjective rating of level 7 from the derived function (Fig. 1B). This individualized intensity was then used in the following Money-Pain tradeoff task.

### Oxytocin/placebo administration

The oxytocin and placebo administration procedure was similar to that used in the previous studies^55,57^. For each treatment, oxytocin or placebo (saline) spray was administered to subjects thrice, consisting of one inhalation of 4 international units (IU) into each nostril resulting a total of 24 IU. The order of oxytocin and placebo sessions was counterbalanced across subjects. After each session, subjects were asked to rate on a 5-point scale about their perception of the administered substance being oxytocin, with 1 indicating it was unlikely to be oxytocin, 5 indicating strong confidence that it was oxytocin. Subjects’ rating indicated that they had no bias towards the substance used in each session (placebo session: mean = 3.13 ± 0.89; oxytocin session: mean = 3.11 ± 1.16 ; the difference between the two sessions: *t*45 = 0.0965, *P* = 0.924). No subjects reported any side effect after the experiments.

### Experimental design

Study 1 adopted a 2 (decision context: gain vs. loss) × 2(shock recipient: self vs. other) within-subject design (Fig.1A). Each subject had to choose between two options representing the tradeoff between money and shock. In the gain context, subjects decided whether to inflict more pain (more electric shocks) either on themselves (gain- self condition) or on the receivers (gain-other condition) to gain more money. In the loss context, subjects would decide whether to inflict more pain either on themselves (loss-self condition) or on the receivers (loss-other condition) to avoid a bigger monetary loss. The sequence of the gain and loss contexts were block designed and counterbalanced across subjects. In study 2, we employed a 2 (decision context: gain vs. loss) × 2 (shock recipient: self vs. other) × 2 (treatment: placebo vs. oxytocin) within-subject design to examine oxytocin’s effect on subjects’ hyperaltruistic preference. Each subject performed the same money-pain tradeoff tasks (same as study 1) twice. Approximately 35 minutes before each experiment and immediately after the pain calibration procedure, each subject was intranasally administered 24 IU oxytocin or placebo (saline) (Fig.1B).

In the Money-Pain tradeoff task, subjects were asked to choose between options associated with different levels of electric shocks and monetary amounts. Crucially, more painful options were always associated with larger monetary gain (or smaller monetary loss), while less painful options offered smaller monetary gains (or larger monetary losses). The experimental task was coded with Psychtoolbox (version 3.0.17) in Matlab (2018b). We first created 60 gain trials using the procedure reported in earlier research^4^. Specifically, each trial was determined by a combination of the differences of shocks (Δs, ranging from 1 to 19, with increment of 1) and money (Δm, ranging from ¥0.2 to ¥19.8, with increment of ¥0.2) between the two options, resulting in a total of 19×99=1881 pairs of [Δs, Δm]. To ensure the trials were suitable for most subjects, we evenly distributed the desired ratio Δm / (Δs + Δm) between 0.01 and 0.99 across 60 trials for each condition. For each trial, we selected the closest [Δs, Δm] pair from the [Δs, Δm] pool to the specific Δm / (Δs + Δm) ratio, which was then used to determine the actual money and shock amounts of two options. The shock amount (*S_less_*) for the less painful option was an integer drawn from the discrete uniform distribution [1∼19], constraint by S_less_ + Δs < 20. Similarly, the money amount (*M_less_*) for the less painful option was drawn from a discrete uniform distribution [¥0.2 ∼ ¥19.8], with the constraint of M_less_ + Δm < 20. Once the *S_less_* and *M_less_* were selected, the shock (*S_more_*) and money (M_more_) magnitudes for the more painful option were calculated as: *S_more_* = S_less_ + Δs, *M_more_* = M_less_ + Δm. For the loss condition, we flipped the signs for the monetary amount and then switched the monetary amounts between the more and less painful options. For example, the two options [¥15 & 10 shocks; ¥10 & 5 shocks] in the gain condition would turn to [- ¥10 & 10 shocks; - ¥15 & 5 shocks] in the loss condition. Subjects completed 60 trials each for all the 4 decision-context and shock-recipient combinations (gain-self, gain-other, loss-self and loss other) thus yielding a total of 240 trials delivered across two blocks (gain and loss blocks). We randomly generated each subject’s trial set using the above method. Across trials, Δs and Δ m was uncorrelated (*r* = -0.0012, *P* = 0.986). The trial sequences and the more/less painful option location (left vs. right) on the computer screen was randomized across subjects within each block.

For each trial, subjects had a maximum of 6s to choose either the more or less painful option by pressing a keyboard button with their right or left index finger. Button presses resulted in the chosen option being highlighted on the screen for 1s. If no choice was entered during the 6s response window, a message ‘Please respond faster!’ was displayed for 1s, and this trial was repeated. An inter-trial interval (ITI) of 1s was introduced before the beginning of the next trial (Fig.1B). Each subject was endowed with ¥40 at the beginning of the task (the maximum amount subjects could have lost in each loss-self and loss-other trials together). At the end of the experiment, one trial for each experimental condition was randomly selected and subjects were renumerated and shocked according to the combined payoff (four conditions) and electric shocks (only in gain-self and loss-self conditions). Each subjects received a ¥60 show up fee in study1(¥220 in study2) and the average total subject fee is ¥97(¥318 in study2).

### Personality trait measures and questionnaires

The Oxford utilitarianism scale was utilized to evaluate two independent dimensions that drive people’s prosocial preference in a moral dilemma^1^: the dimension of instrumental harm (IH) measures individuals’ willingness to compromise moral principles by either breaking rules or causing harm to others to achieve favorable outcomes; whereas the impartial beneficence (IB) captures their levels of impartial concern for the well-being of the general population. IH has four items including statements such as “Sometimes it is morally necessary for innocent people to die as collateral damage if more people are saved overall”. The IB measure contains 5 items such as “It is morally wrong to keep money that one doesn’t really need if one can donate it to causes that provide effective help to those who will benefit a great deal”. Subjects responded to each IH and IB item via a 7-point Likert scale. IH and IB scores were derived by calculating the average score of all items on the subscale for each subject. We also measured subjects’ empathetic concern (EC), the identification with the whole of humanity and concern for future generation, using the interpersonal reactivity index (IRI) ^78^. The EC comprises 7 items with a 5-point rating scale and has been shown to be correlated with both IH and IB (see Supplementary Fig. 2). Therefore, we performed multiple regression analysis to examine the relationships between IB, IH and moral behaviors, with EC included as a covariate of no interest.

All subjects completed debriefing questionnaires that assessed their beliefs about the experimental setup, such as whether they believed that the recipient would actually receive the electric shocks. Moreover, for study 2, we also collected another questionnaire by asking subjects how they perceived the task structure (whether they regarded the experiment as helping or harming other subjects) in both decision contexts (gain and loss). Subjects rated their moral perception within gain and loss contexts using a scale ranging from -4 to 4. Positive values indicated harm frame (inflicting pain on the receiver to increase monetary gain/avoid monetary loss), whereas negative values indicated help frame (sacrificing greater monetary gain/accepting greater monetary loss to alleviate others’ pain) with 0 indicating neutrality. Therefore, a larger rating indicated subject’s moral perception of a more harming frame.

### Data Analysis

#### Harm aversion model

We analyzed subjects’ choice data using the harm aversion model^6^, where choices were driven by the subjective value difference (ΔV) between the less and more painful options (Eq. 1). Parameter κ (0 ≤ κ ≤ 1) quantifies the relative weight deciders attribute to electric shock versus money.

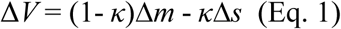

We separately estimated κ in the four conditions related to both the shock recipients (self vs. other) and decision contexts (gain vs. loss) and thus yielded *κ_gain-self_*, *κ_gain-other_*, *κ_loss-self_*, and *κ_loss-other_* respectively. Δm and Δs denoted the objective differences in money and electric shocks between the more and less painful options. In this model, trial-wise ΔV was transformed into choice probability via a softmax function where γ serves as a subject-specific parameter for choice consistency (Eq. 2) (see Supplementary Fig. 10 for the model comparison results that confirmed a single γ for all conditions fitted subjects’ behavior better than other candidate models).

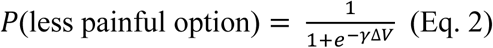

Model parameters were estimated for each subject using maximum likelihood estimation (MLE), and the maximization was achieved using the fminsearchbnd function in MATLAB (Mathworks). We repeated the estimation with 300 random initial parameter values to achieve stable parameter estimators.

#### Regression models

We applied the mixed-effect logistic regression model to investigate how Δm and Δs independently influenced subjects’ choices. Although this regression model seems equivalent to the harm aversion model mentioned above, the regression coefficients obtained from the regression model allowed us to separately examine how the decision contexts and oxytocin treatment affected subjects’ sensitivities towards Δm and Δs. In the regression model 1 of Study 1, choices were coded as 1 if subjects chose the less painful option and 0 otherwise. Additionally, decision context (1 for gain and 0 for loss) and shock recipient (1 for self and 0 for other) were treated as categorical variables. Each regressor had a fixed effect across all subjects and a random effect specific to each subject. The regression model 1 was specified as following^79^:

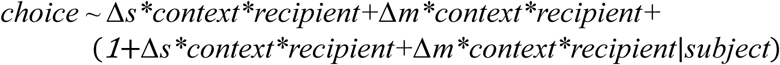

To examine how oxytocin affected subjects’ behavior, in Study 2, we expanded regression model 1 by including the variable of treatment (1 for oxytocin treatment and 0 for placebo) as an additional categorical independent variable. The regression model 2 was described as:

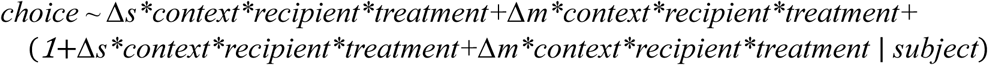

#### Moderated mediation analysis

We set out to examine whether subjects’ internal moral framing mediates the association between their personality traits (IH) and moral preferences, and more importantly whether the mediation is context (gain vs. loss) specific. To test the moderated mediation, we constructed a mediation model where the mediator (M):

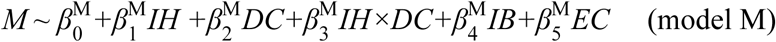

and the dependent variable model was:

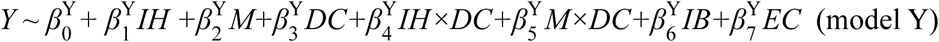

Where Y, M respectively referred to the dependent variable (relative harm sensitivity) and the mediator (harm framing report). Variables IH, DC, IB and EC corresponded to independent variable instrumental harm attitude (IH), decision context (gain vs. loss, the moderator variable), impartial beneficence (IB, variable of no interest) and empathic concern (EC, variable of no interest), accordingly. The existence of interaction effect (*β_3_^M^*,*β_4_^Y^*, *β_5_^Y^*) in either path a (X→M), path b (M→Y) or path c’ (X→Y)in the model is sufficient to claim moderation of mediation^80^.

We utilized the bootstrapping procedure with 5000 bootstrap resamples to analyzed the data^81,82^. This analysis was conducted separately for the placebo and oxytocin sessions. Our results showed that the interaction coefficients (placebo: *β_4_*^Y^= 0.036 ± 0.029, *t*_84_ = 1.267, *P* = 0.209; *β_5_*^Y^ = 0.0242 ± 0.017, *t*_84_ = 1.415, *P* = 0.161; Oxytocin: *β_4_*^Y^ = 0.006 ± 0.010, *t*_84_ = 0.319, *P* = 0.751 ; *β_5_*^Y^= 0.003 ± 0.012, *t*_84_=0.246, *P* = 0.806) in model Y were not significant, while *β*^M^ was significant in the placebo session (*β*^M^= 0.784 ± 0.343, *t*86 = 2.288, *P* = 0.025) but not the oxytocin session (*β*^M^= -0.137 ± 0.291, *t*86 = -0.472, *P* = 0.638, indicating that the moderation effect present only on the path a (see Fig. 6B) in the placebo condition. This aligns with our findings that decision contexts modulated the correlation between IH and relative harm sensitivities (Fig. 3C & Fig. 5C). Therefore, we report all the results from the reduced version of the moderated mediation model (model Y^5^, see supplementary Table S3 for details).

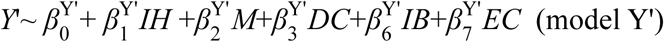

#### Statistics and software

ANOVAs and non-parametric analyses were conducted with SPSS 27.0, whereas regression analyses were performed using “fitglme” and “fitlm” functions in Matlab (2022b). The moderated mediation analysis was performed using “bruceR” and “mediation” packages in R (version 4.2.2). All the reported p-values are two-tailed.

## Data availability

The data and analysis code that support the findings of this study are available at https://osf.io/tpfeg/.

## Acknowledgement

This work was supported by the National Science and Technology Innovation 2030 Major Program (2021ZD0203702), National Natural Science Foundation of China Grants (32071090) to J.L.

---

**Supplementary Fig. 1.**
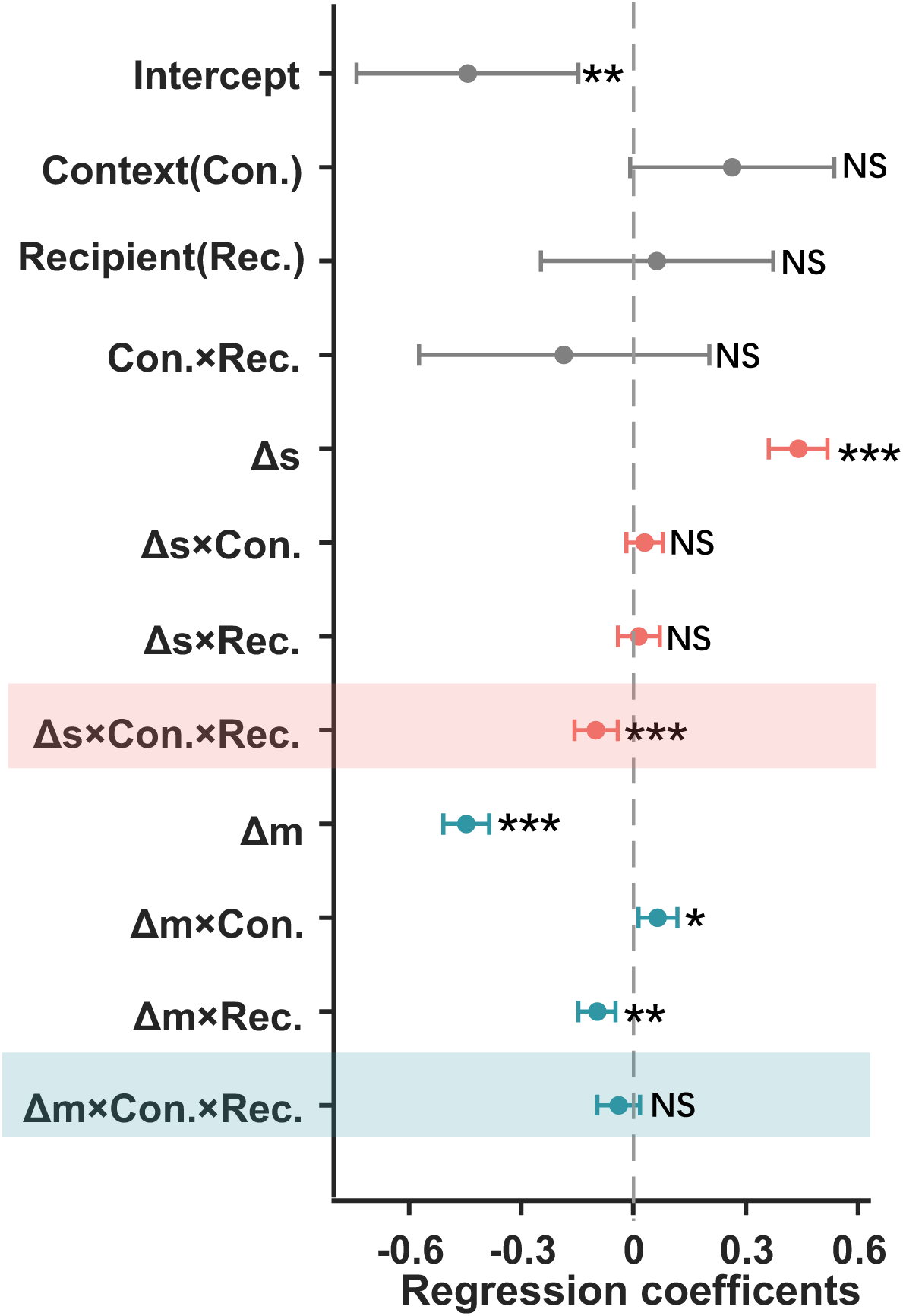
Mixed-effect logistic regression analysis results for experiment 1(regression model 1). Δm and Δs represented the objective differences in money and electric shocks between the more and less painful options. Δ*s* and Δ*m* are numerical variables, while the context (gain vs. loss) and recipient (other vs. self) are categorial variables. The Δs × context × recipient interaction and Δm × context × recipient interaction was further elucidated in Fig. 2C and D. Con: context; Rec: recipient. Error bars represent 95% confidence interval (CI). NS, not significant, * *P* < 0.05, ** *P* < 0.01 and *** *P* < 0.001.

**Supplementary Fig. 2.**
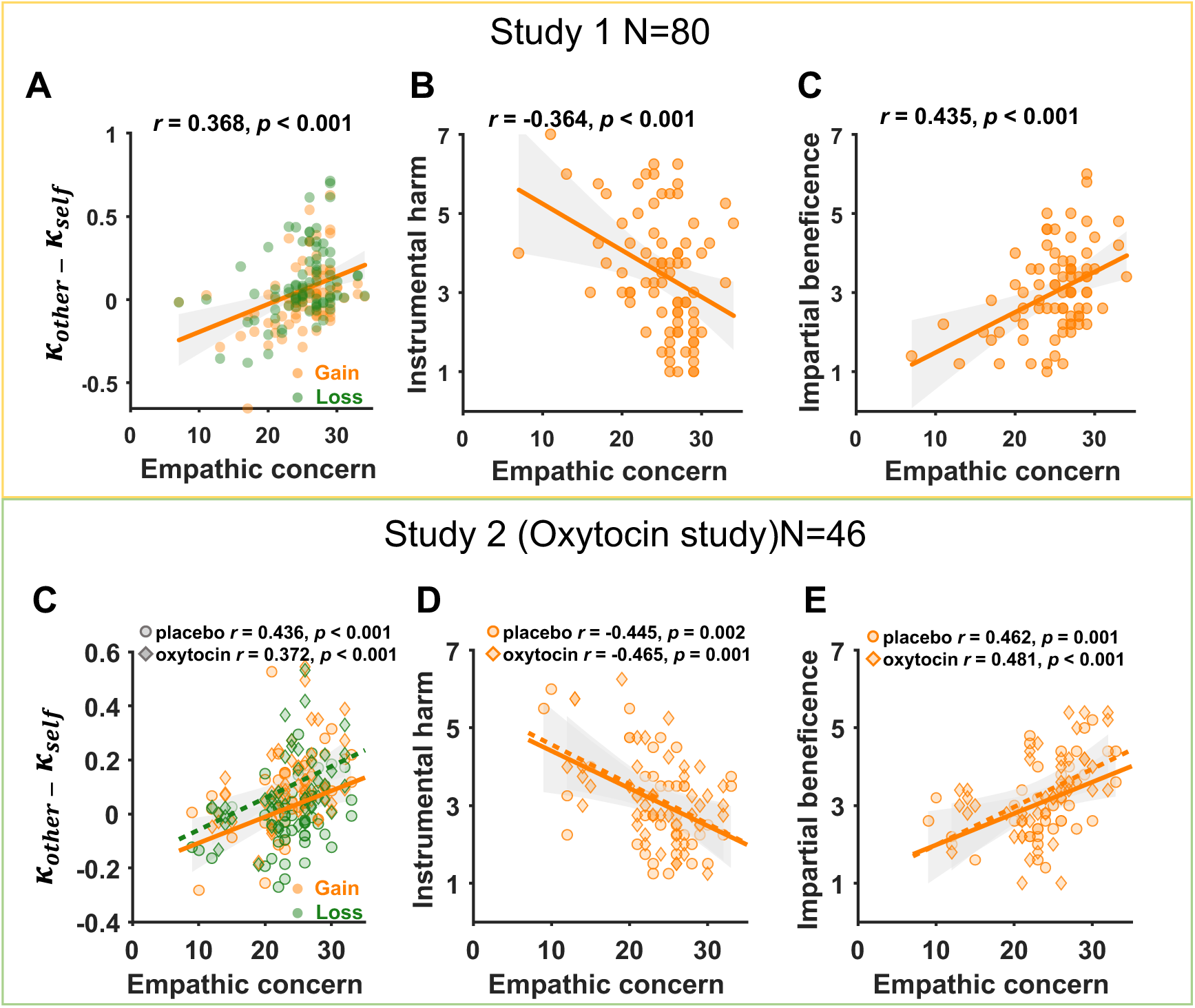
The relationship between personality trait empathic concern (EC) and utilitarian moral personality traits. Correlation between EC and hyperaltruism (*κ_other_* – κ*_self_*), instrumental harm (IH), impartial beneficence (IB) both in the Study1(**A-C**) and Study 2 (**D-E**). All the correlations were performed using Pearson correlations.

**Supplementary Fig. 3.**
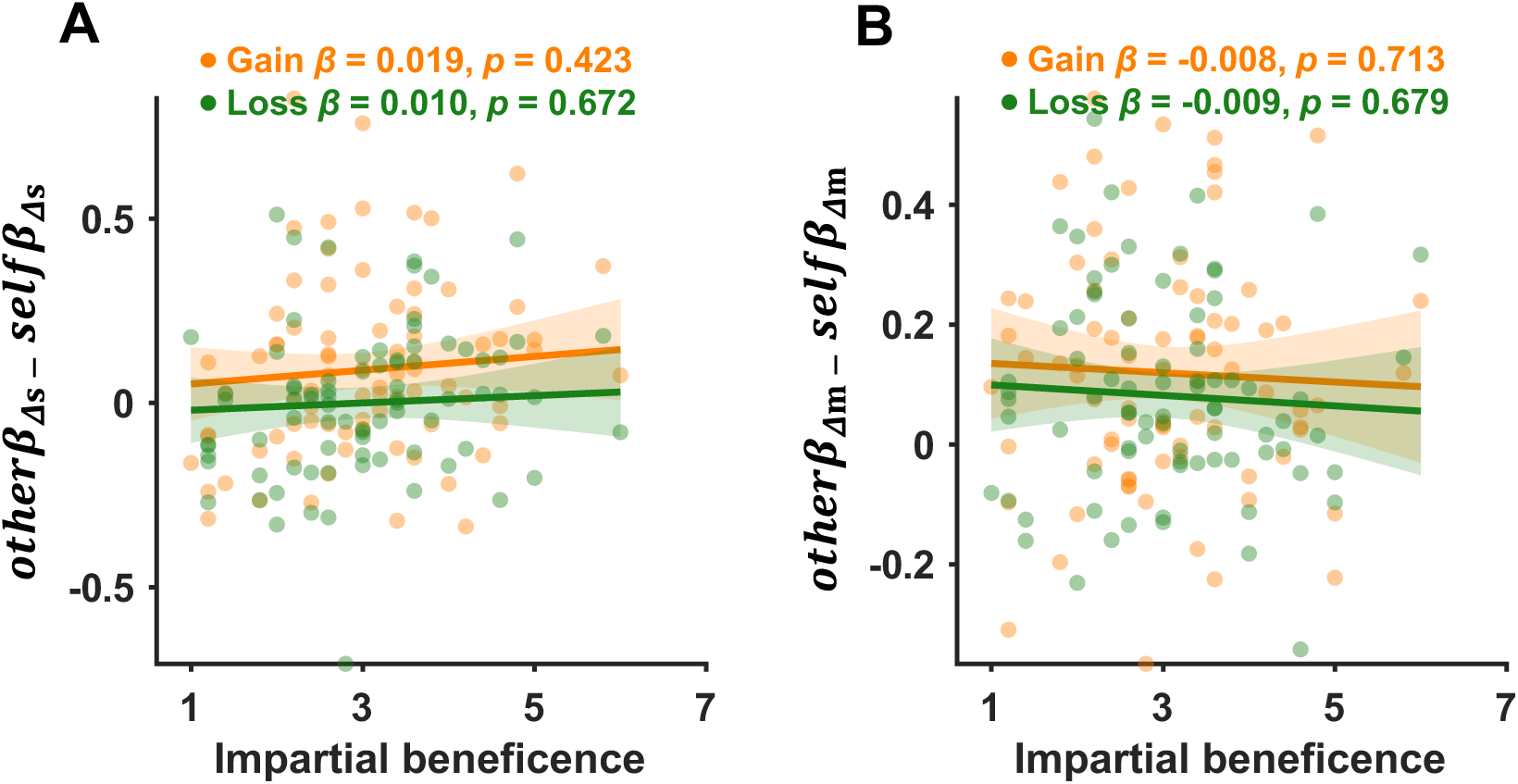
Association between impartial beneficence (IB) and subjects’ relative harm/money sensitivities in Study 1. IB showed no significant correlation with subjects’ relative harm sensitivities (*otherβ_Δs_* − *selfβ_Δs_*) **(A)** or relative monetary sensitivities (*otherβ_Δm_* − *selfβ_Δm_*) **(B)**. The regression coefficients showed the association between IB and relative harm/monetary sensitivities, controlling for empathic concern and instrumental harm.

**Supplementary Fig. 4.**
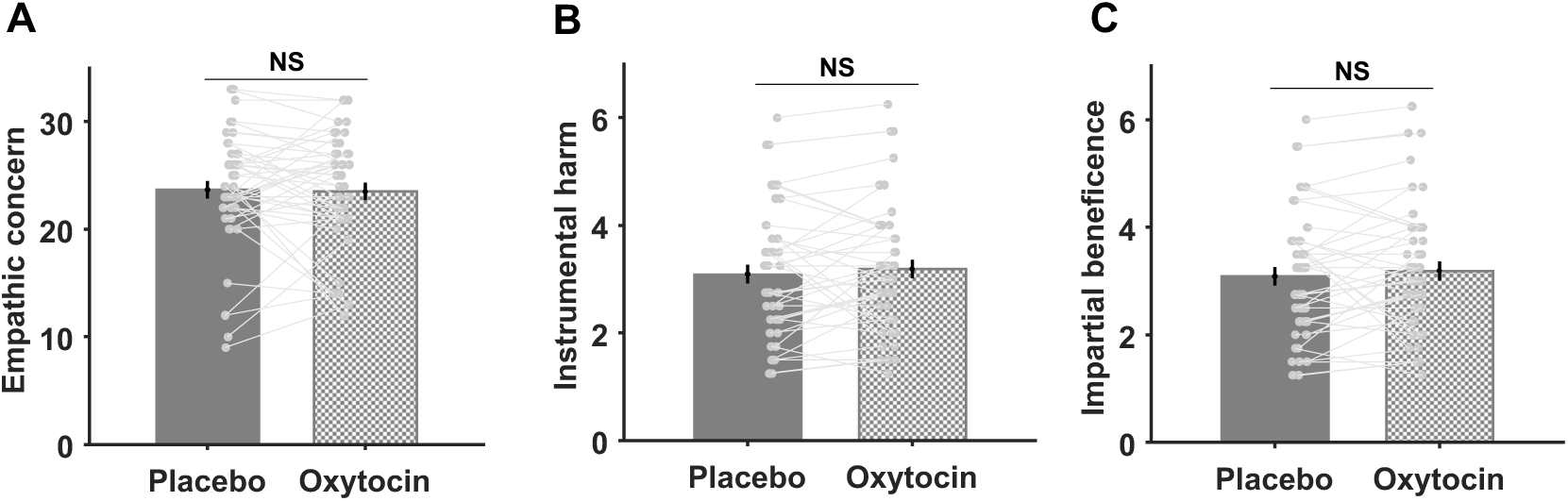
Oxytocin did not influence personality traits in study 2. (**A-C**) Oxytocin did not affect subjects’ ratings on empathic concern (EC), instrumental harm (IH) or impartial beneficence (IB). Error bars represent SE. NS, not significant.

**Supplementary Fig. 5.**
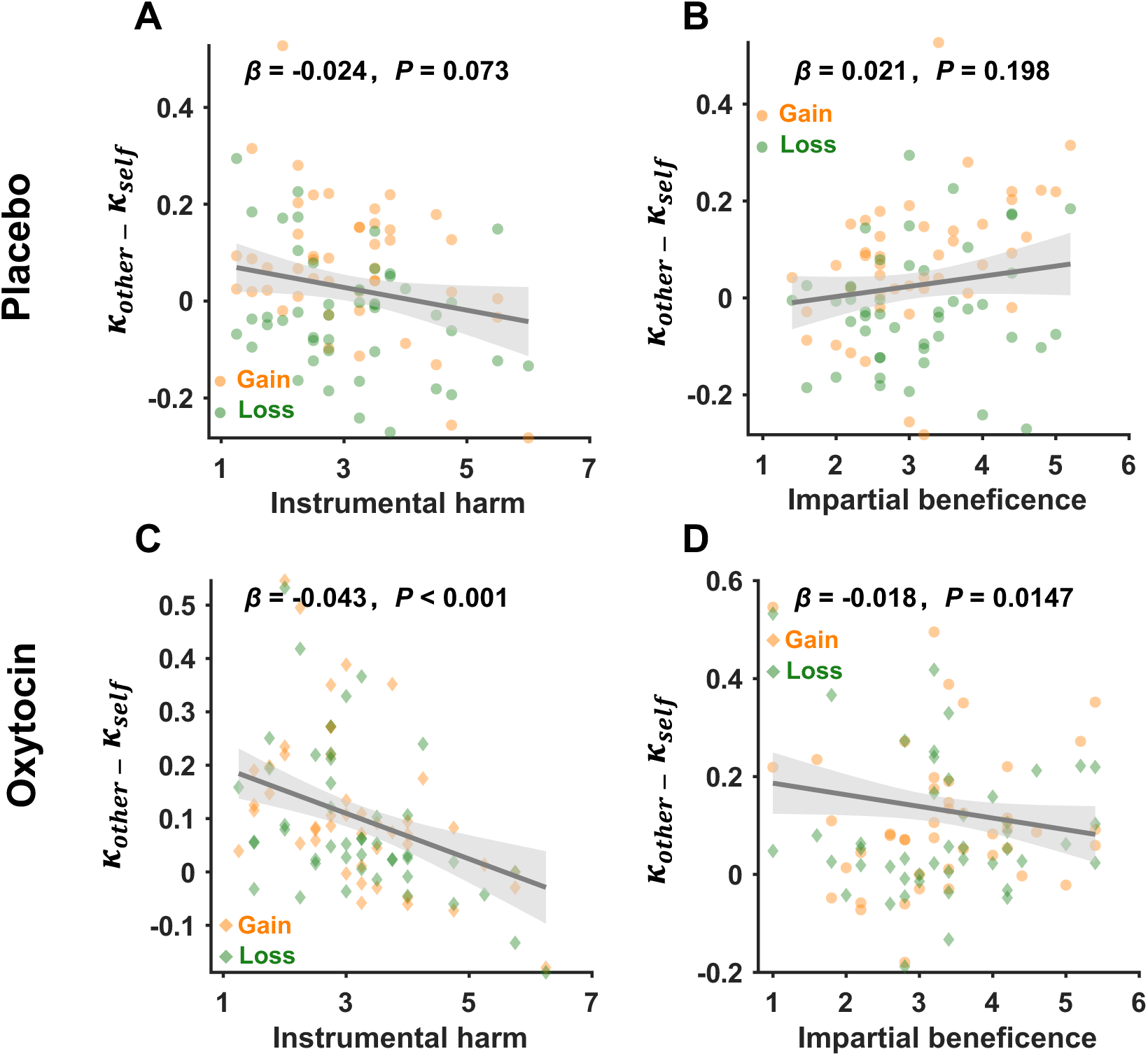
Relationships between hyperaltruism and utilitarian moral personality traits in Study 2. (**A-B**) In the placebo session, hyperaltruistic preferences showed a marginally negative correlation with instrumental harm (IH) but no significant correlation with impartial beneficence (IB). (**C-D**) In the oxytocin session, hyperaltruistic preferences exhibited significantly negative correlation with IH yet still no significant correlation with IB. The regression coefficients in Figure A, C showed the relationship between IH and moral preference, controlling for empathic concern (EC) and IB, while the coefficient in Figure B, D showed the relationship between IB and moral preference, controlling for EC and IH.

**Supplementary Fig. 6.**
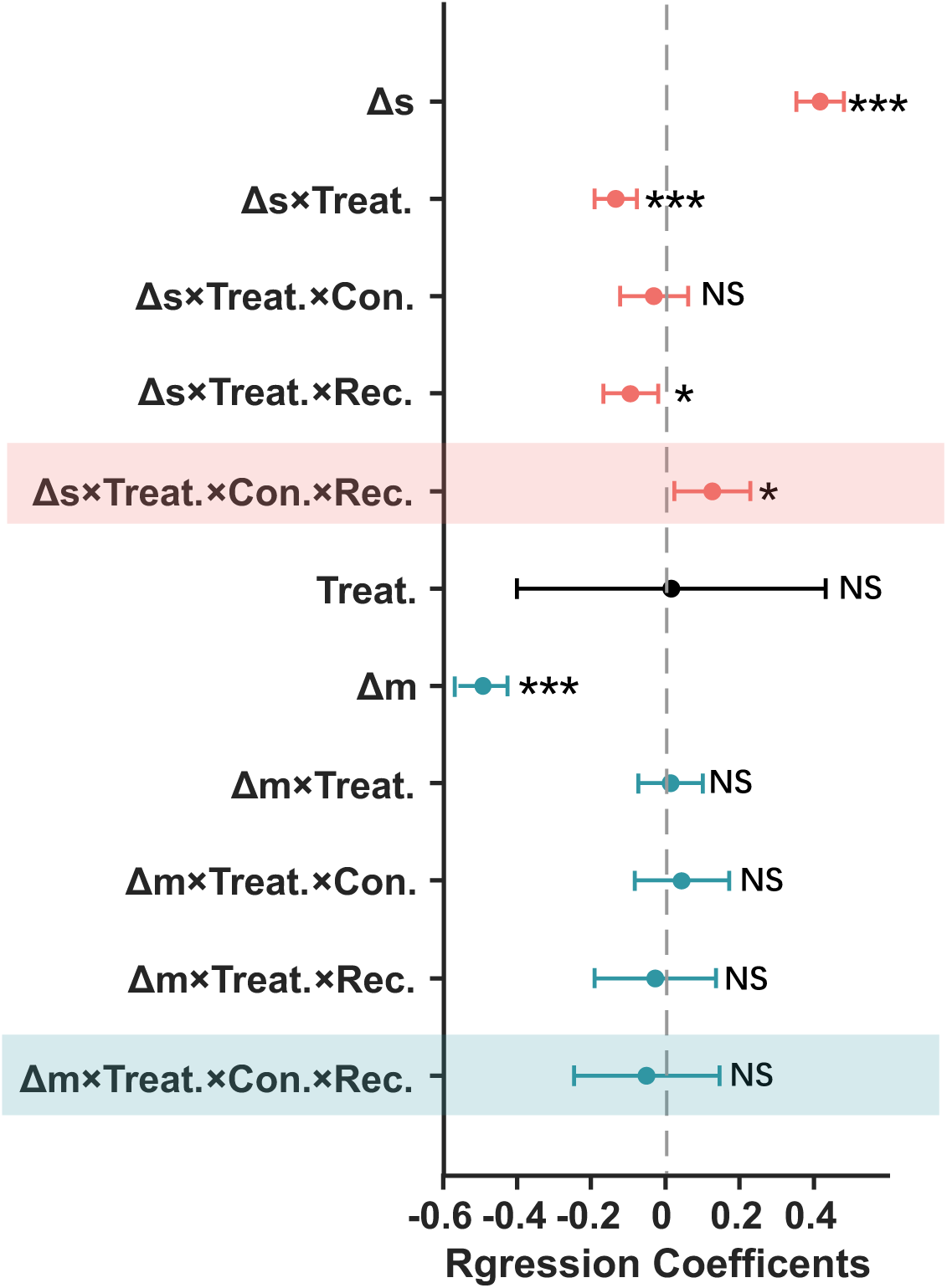
The main results of mixed-effect logistic regression analysis in study 2(regression model 2). Δm and Δs represented the objective differences in money and electric shocks between the more and less painful options. The Δs × Treat.× Con.× Rec. interaction was further explained in Fig. 5A and Supplementary Fig.8. The Δm × Treat.× Con.× Rec. interaction was further explained in Fig. 5B. Treat.: treatment (placebo vs. oxytocin); Con.: decision contexts (gain vs. loss), and Rec.: recipient (self vs. other) are all categorial variables. Error bars represent 95% confidence interval (CI). NS, not significant, * *P* < 0.05, and *** *P* < 0.001.

**Supplementary Fig. 7.**
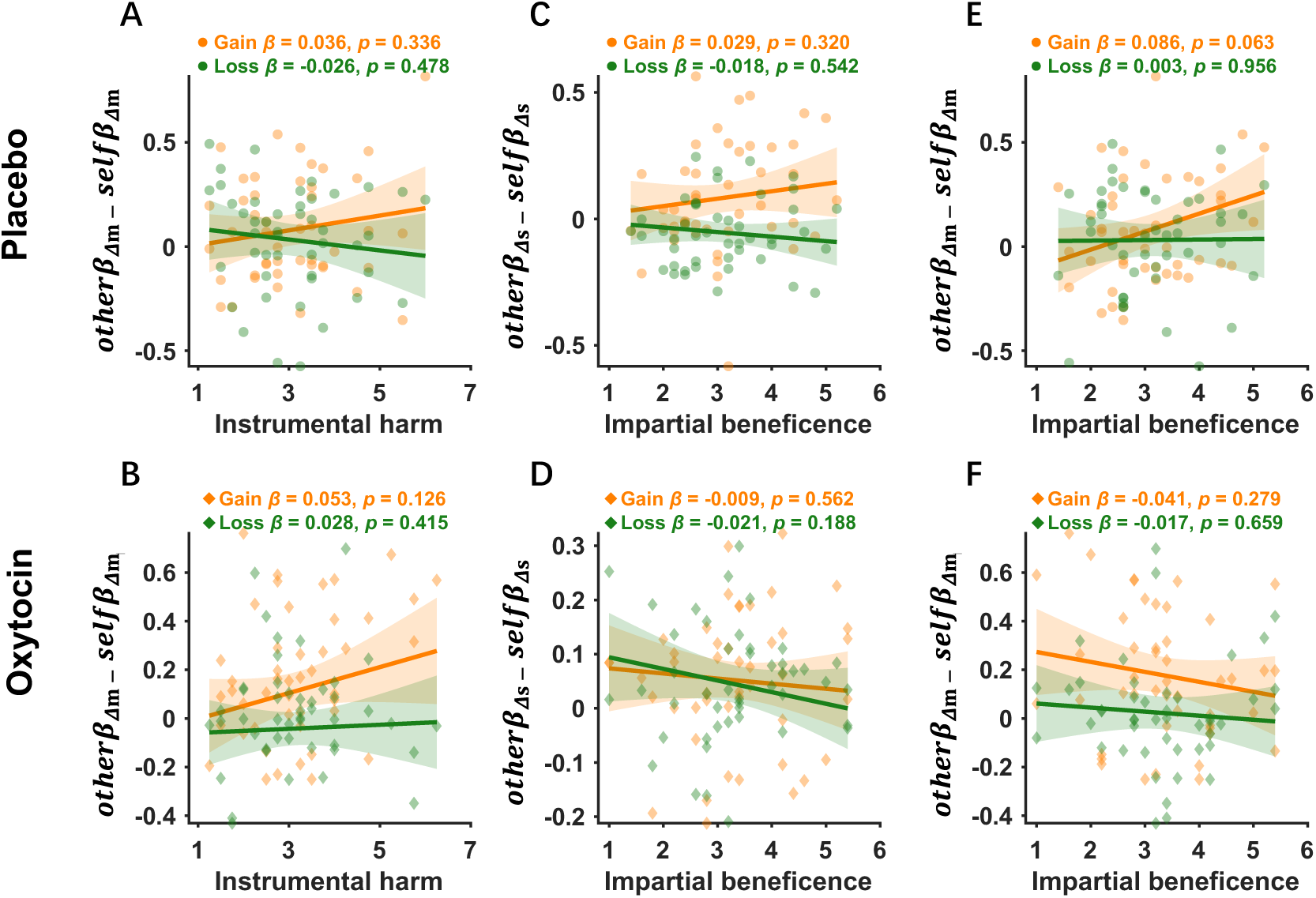
Associations between relative harm/money sensitivities and utilitarian moral personality traits in Study 2. (**A-B**) The relative monetary sensitivities were not significantly related with instrumental harm (IH) in either placebo or oxytocin condition. (**C-D**) In both the placebo and oxytocin conditions, there was no significant correlation between relative harm sensitivity and impartial beneficence (IB). (**E-F**) No significant correlation was found between relative monetary sensitivities and IB in either the placebo or oxytocin conditions. Figure A & B illustrated the relationship between IH and relative monetary sensitivities, with empathic concern (EC) and IB controlled. Figure C ∼ F described the relationship between IB and relative harm/monetary sensitivities, controlling for EC and IH.

**Supplementary Fig. 8.**
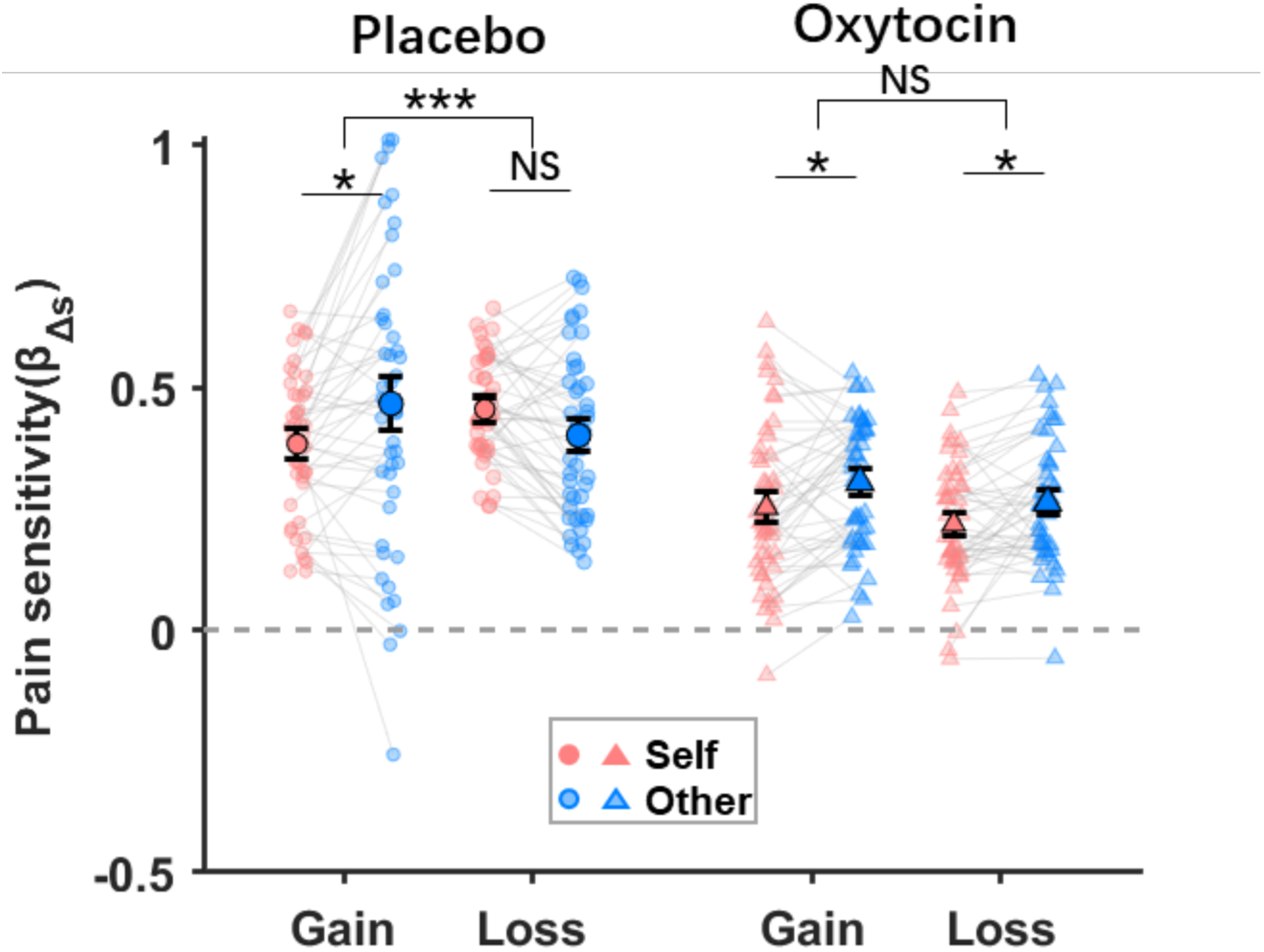
Pain sensitivity across experimental conditions. The administration of oxytocin significantly reduced participants’ pain sensitivity, yet also restored the harm sensitivity patterns in both the gain and loss conditions relative to the placebo treatment. Error bars represent SE across subjects. NS, not significant, * *P* < 0.05, and *** *P* < 0.001.

**Supplementary Fig. 9.**
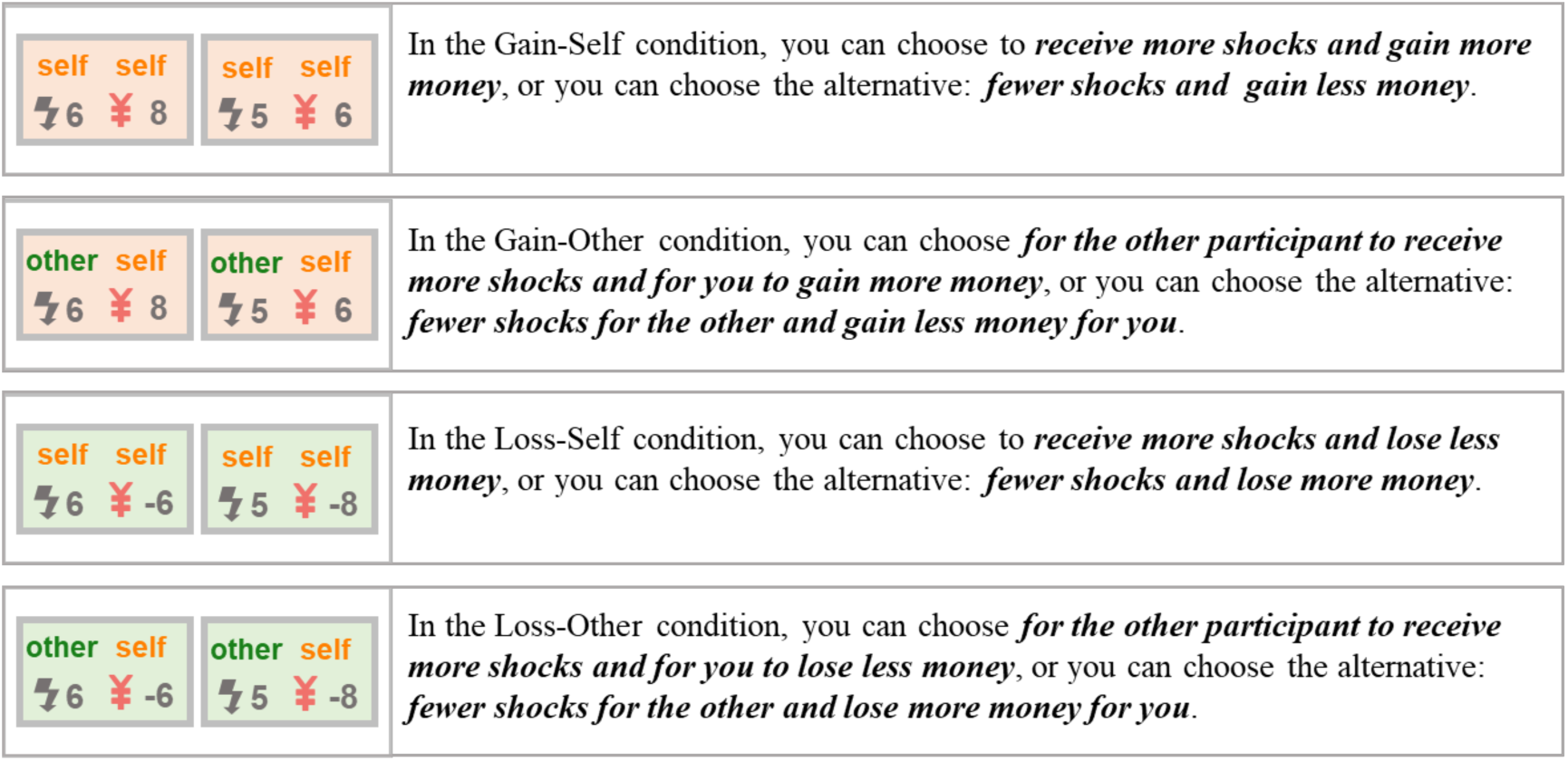
The experimental instructions provided to subjects. We presented the options to subjects in a neutral and descriptive manner, focusing only on the relevant components (shocks and money) across four conditions.

**Supplementary Fig. 10.**
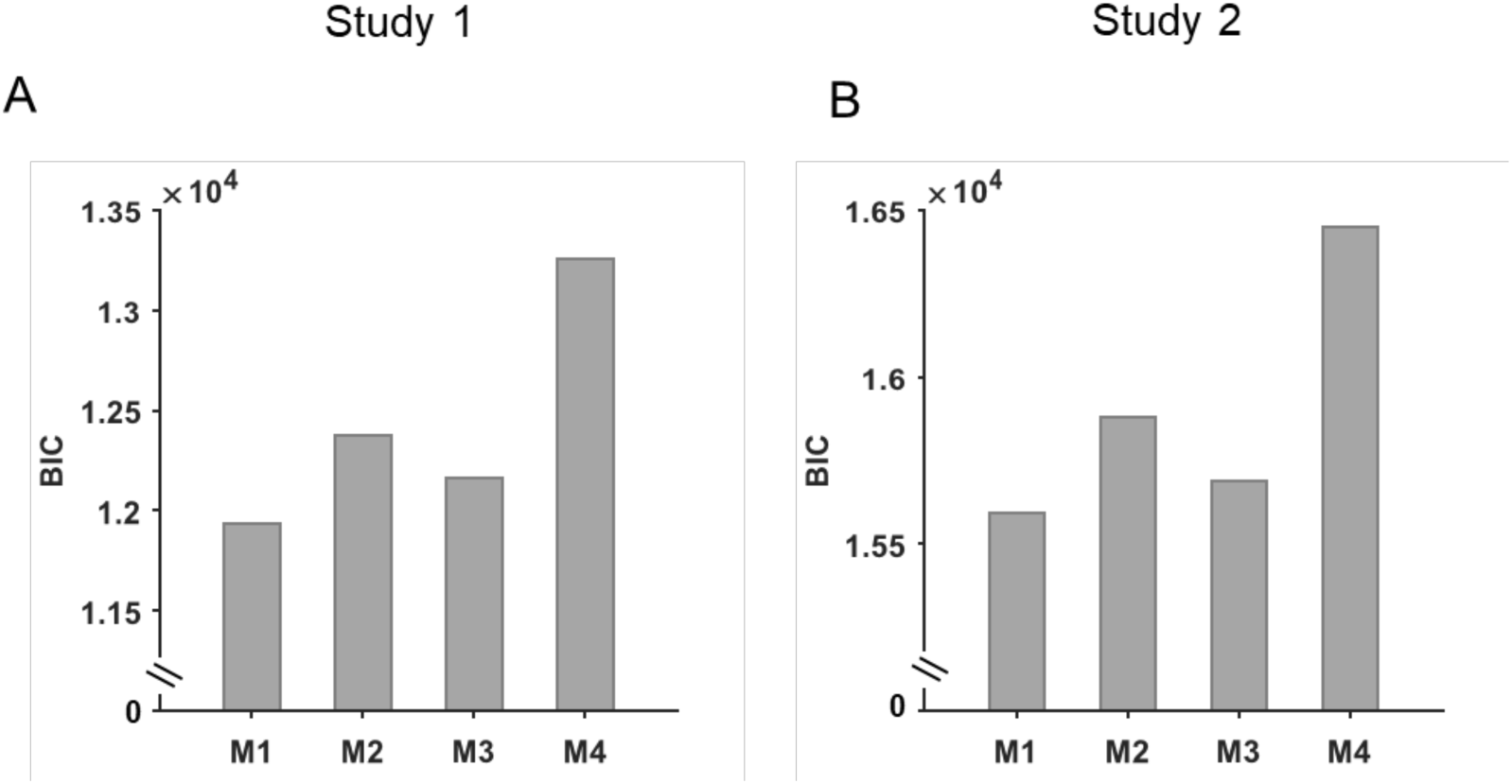
Model comparison results. To investigate whether the choice consistency parameter (γ) is condition specific, we compared four candidate models: (M1) γ was constant across all conditions; (M2) γ varied based on the pain recipient (self vs. other); (M3) γ varied between decision contexts (gain vs. loss); and (M4) γ varied across all four conditions (self-gain, other-gain, self-loss, and other-loss). Model 1 (M1) performed the best among all the four candidate models both in Study 1 **(A)** and Study 2 **(B)**.

**Table S1.**
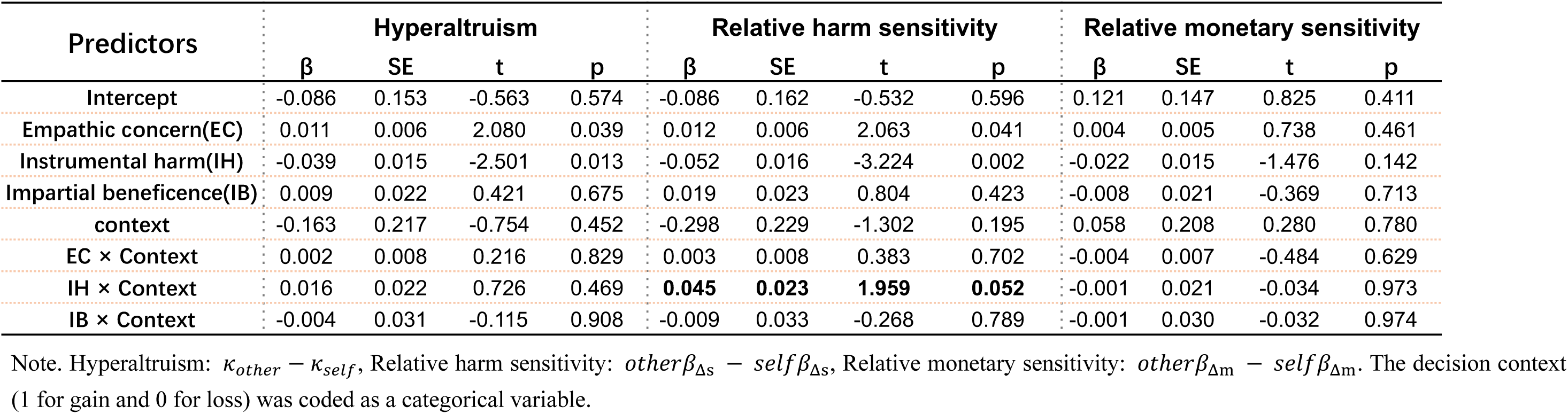
Contribution of EC, IH and IB scores on subjects’ behavior in Study 1.

| Predictors | Hyperaltruism |  |  |  | Relative harm sensitivity |  |  |  | Relative monetary sensitivity |  |  |  |
| --- | --- | --- | --- | --- | --- | --- | --- | --- | --- | --- | --- | --- |
| | $\beta$ | SE | t | p | $\beta$ | SE | t | p | $\beta$ | SE | t | p |
| Intercept | -0.086 | 0.153 | -0.563 | 0.574 | -0.086 | 0.162 | -0.532 | 0.596 | 0.121 | 0.147 | 0.825 | 0.411 |
| Empathic concern(EC) | 0.011 | 0.006 | 2.080 | 0.039 | 0.012 | 0.006 | 2.063 | 0.041 | 0.004 | 0.005 | 0.738 | 0.461 |
| Instrumental harm(IH) | -0.039 | 0.015 | -2.501 | 0.013 | -0.052 | 0.016 | -3.224 | 0.002 | -0.022 | 0.015 | -1.476 | 0.142 |
| Impartial beneficence(IB) | 0.009 | 0.022 | 0.421 | 0.675 | 0.019 | 0.023 | 0.804 | 0.423 | -0.008 | 0.021 | -0.369 | 0.713 |
| context | -0.163 | 0.217 | -0.754 | 0.452 | -0.298 | 0.229 | -1.302 | 0.195 | 0.058 | 0.208 | 0.280 | 0.780 |
| EC × Context | 0.002 | 0.008 | 0.216 | 0.829 | 0.003 | 0.008 | 0.383 | 0.702 | -0.004 | 0.007 | -0.484 | 0.629 |
| IH × Context | 0.016 | 0.022 | 0.726 | 0.469 | **0.045** | **0.023** | **1.959** | **0.052** | -0.001 | 0.021 | -0.034 | 0.973 |
| IB × Context | -0.004 | 0.031 | -0.115 | 0.908 | -0.009 | 0.033 | -0.268 | 0.789 | -0.001 | 0.030 | -0.032 | 0.974 |

Note. Hyperaltruism: $\kappa_{other} - \kappa_{self}$ , Relative harm sensitivity: $other\beta_{\Delta s} - self\beta_{\Delta s}$ , Relative monetary sensitivity: $other\beta_{\Delta m} - self\beta_{\Delta m}$ . The decision context (1 for gain and 0 for loss) was coded as a categorical variable.

**Table S2.** Contribution of EC, IH and IB scores on subjects’ behavior in Study 2.

| Predictors | Hyperaltruism |  |  |  | Relative harm sensitivity |  |  |  | Relative monetary sensitivity |  |  |  |
| --- | --- | --- | --- | --- | --- | --- | --- | --- | --- | --- | --- | --- |
| | $\beta$ | SE | t | p | $\beta$ | SE | t | p | $\beta$ | SE | t | p |
| Intercept | -0.085 | 0.123 | -0.688 | 0.493 | 0.065 | 0.170 | 0.382 | 0.704 | -0.205 | 0.266 | -0.771 | 0.443 |
| Empathic concern(EC) | 0.005 | 0.004 | 1.162 | 0.248 | 0.006 | 0.006 | 1.086 | 0.281 | -0.004 | 0.009 | -0.444 | 0.658 |
| Instrumental harm(IH) | -0.032 | 0.017 | -1.904 | 0.060 | -0.071 | 0.024 | -3.005 | 0.003 | 0.035 | 0.037 | 0.967 | 0.336 |
| Impartial beneficence(IB) | 0.047 | 0.021 | 2.200 | 0.031 | 0.029 | 0.029 | 1.000 | 0.320 | 0.086 | 0.046 | 1.887 | 0.063 |
| context | -0.034 | 0.174 | -0.197 | 0.844 | -0.340 | 0.241 | -1.412 | 0.162 | 0.323 | 0.375 | 0.862 | 0.391 |
| EC × Context | 0.002 | 0.006 | 0.332 | 0.740 | 0.005 | 0.008 | 0.601 | 0.549 | 0.003 | 0.012 | 0.264 | 0.792 |
| IH × Context | 0.018 | 0.024 | 0.737 | 0.463 | **0.077** | **0.033** | **2.297** | **0.024** | -0.062 | 0.052 | -1.187 | 0.238 |
| IB × Context | -0.051 | 0.030 | -1.711 | 0.091 | -0.047 | 0.042 | -1.140 | 0.257 | -0.084 | 0.065 | -1.295 | 0.199 |

b. Oxytocin condition
| Predictors | Hyperaltruism |  |  |  | Relative harm sensitivity |  |  |  | Relative monetary sensitivity |  |  |  |
| --- | --- | --- | --- | --- | --- | --- | --- | --- | --- | --- | --- | --- |
| | $\beta$ | SE | t | p | $\beta$ | SE | t | p | $\beta$ | SE | t | p |
| Intercept | 0.138 | 0.125 | 1.100 | 0.275 | 0.056 | 0.109 | 0.511 | 0.611 | 0.049 | 0.253 | 0.195 | 0.846 |
| Empathic concern(EC) | 0.010 | 0.004 | 2.397 | 0.019 | 0.006 | 0.004 | 1.675 | 0.098 | 0.001 | 0.009 | 0.157 | 0.876 |
| Instrumental harm(IH) | -0.047 | 0.017 | -2.721 | 0.008 | -0.037 | 0.015 | -2.458 | 0.016 | 0.053 | 0.035 | 1.541 | 0.126 |
| Impartial beneficence(IB) | -0.034 | 0.019 | -1.813 | 0.073 | -0.009 | 0.016 | -0.582 | 0.562 | -0.041 | 0.038 | -1.091 | 0.279 |
| Context(gain vs. loss) | -0.071 | 0.177 | -0.401 | 0.690 | 0.092 | 0.154 | 0.596 | 0.553 | -0.372 | 0.358 | -1.040 | 0.301 |
| EC × Context | 0.002 | 0.006 | 0.385 | 0.701 | -0.002 | 0.005 | -0.336 | 0.737 | 0.012 | 0.012 | 0.985 | 0.327 |
| IH × Context | 0.008 | 0.024 | 0.323 | 0.747 | **-0.006** | **0.021** | **-0.272** | **0.786** | -0.045 | 0.049 | -0.918 | 0.361 |
| IB × Context | -0.010 | 0.026 | -0.384 | 0.702 | -0.012 | 0.023 | -0.526 | 0.600 | 0.024 | 0.053 | 0.457 | 0.649 |

Note. Hyperaltruism: $\kappa_{other} - \kappa_{self}$ , Relative harm sensitivity: $other\beta_{\Delta s} - self\beta_{\Delta s}$ , Relative monetary sensitivity: $other\beta_{\Delta m} - self\beta_{\Delta m}$ . The decision context (1 for gain and 0 for loss) was coded as a categorical variable.

**Table S3.** Moderated mediation analysis results for Study 2.

| Predictors | Placebo |  | Oxytocin |  |
| --- | --- | --- | --- | --- |
|  | Mediator | Dependent variable | Mediator | Dependent variable |
| | Harm framing report<br>$\beta$(SE) | Relative harm sensitivity<br>$\beta$(SE) | Harm framing report<br>$\beta$(SE) | Relative harm sensitivity<br>$\beta$(SE) |
| Intercept | 1.196(0.279)*** | 0.069(0.023)** | 1.609(0.239)*** | 0.049(0.014)*** |
| Context | -0.739(0.396) | -0.108(0.032)** | -0.196(0.338) | -0.002(0.019) |
| Empathic concern(EC) | 0.020(0.046) | 0.008(0.004)* | 0.023(0.040) | 0.005(0.002)* |
| Impartial beneficence(IB) | -0.116(0.239) | 0.010(0.019) | -0.270(0.179) | -0.008(0.010) |
| Instrumental harm(IH) | **-0.758(0.257)\*\*\***<br>**a** | **-0.018(0.016)**<br>**c'** | **-0.753(0.220)\*\*\***<br>**a** | **-0.017(0.011)**<br>**c'** |
| Context × IH | **0.784(0.343)\*** | — | **-0.137(0.291)** | — |
| Harm framing report | — | **0.039(0.008)\*\*\***<br>**b** | — | **0.028(0.006)\*\*\***<br>**b** |

Note: In the moderated mediation model, IH serves as the independent variable, harm framing report as the mediator, relative harm sensitivity as the dependent variable, with decision context (gain vs. loss) as the moderator. **a** represents the influence of the independent variable on the mediator, while **b** represents the effect of the mediator on the dependent variable, and **c'** represents the direct effect (insignificant indicating a full mediation effect). The coefficients highlighted in red indicate the moderation effect of the context (gain vs. loss). \* $P < 0.05$ , \*\* $P < 0.01$ and \*\*\* $P < 0.001$ .

## Notes

https://osf.io/tpfeg/

### Competing Interest Statement

The authors have declared no competing interest.

### Summary of Updates

Supplementary figures 8, 9, 10 added to clarify methods and results used; main text revised to better reflect the advances in the field

